# The Role of Biofilms, Bacterial Phenotypes, and Innate Immune Response in *Mycobacterium avium* Colonization to Infection

**DOI:** 10.1101/2021.07.26.453817

**Authors:** Catherine Weathered, Kelly Pennington, Patricio Escalante, Elsje Pienaar

## Abstract

*Mycobacterium avium* complex (MAC), is known for colonizing and infecting humans following inhalation of the bacteria. MAC pulmonary disease is notoriously difficult to treat and prone to recurrence. Both the incidence and prevalence MAC pulmonary disease have been increasing globally.

MAC is well known to form biofilms in the environment, and *in vitro*, these biofilms have been shown to aid MAC in epithelial cell invasion, protect MAC from phagocytosis, and cause premature apoptosis in macrophages. *In vivo*, the system of interactions between MAC, biofilms and host macrophages is complex, difficult to replicate *in vitro* and in animal models, has not been fully characterized. Here we present a three-dimensional agent-based model of a lung airway to help understand how these interactions evolve in the first 14 days post-bacterial inhalation. We parameterized the model using published data and performed uncertainty analysis to characterize outcomes and parameters’ effects on those outcomes.

Model results show diverse outcomes, including wide ranges of macrophage recruitment levels, and bacterial loads and phenotype distribution. Though most bacteria are phagocytosed by macrophages and remain intracellular, there are also many simulations in which extracellular bacteria continue to drive the colonization and infection. Initial parameters dictating host immune levels, bacterial loads introduced to the airway, and biofilm conditions have significant and lasting impacts on the course of these results. Additionally, though macrophage recruitment is key for suppressing bacterial loads, there is evidence of significant excess recruitment that fail to impact bacterial numbers. These results highlight a need and identify a path for further exploration into the inhalation events in MAC infection. Early infection dynamics could have lasting impacts on the development of nodular bronchiectatic or fibrocavitary disease as well as inform possible preventative and treatment intervention targeting biofilm-macrophage interactions.

## 1. Introduction

Non-tuberculous mycobacterial pulmonary disease (NTM-PD) is a growing threat, with both incidence and prevalence steadily rising worldwide^1^. Estimated incidence ranges from 4.1-14.1 per 100,000 people in the US^2^. In many developed countries, NTM-PD cases outnumber those of tuberculosis, a related infection^3^. NTM-PD is notoriously difficult to treat with a success rate between 45-65% even after twelve to twenty-four months of intense antibiotic treatment^4^.

NTMs are prevalent in the environment where they are known to form biofilms. In the environment^5,6^ and many biomedical devices^7,8^, biofilms allow various species of bacteria to communicate and cooperate, share resources, and protect themselves from antimicrobials^9^. Mycobacterial biofilms share some or all of these functions^10^. Recently, NTM biofilms have gotten more attention for their role in establishing and prolonging infections^10^, and *Mycobacterium abscessus* biofilms have been observed in patient lungs^11–13^.

*Mycobacterium avium* complex (MAC) is a member of the NTM family and most common cause of NTM-PD^14^. MAC is known to have multiple phenotypes of bacteria, which live together in colonies, each serving different roles^15^. Planktonic bacteria are faster growing^16^, but more susceptible to antibiotics^17^ and not biofilm-associated. Sessile bacteria grow more slowly^16^ and create biofilms^15^. These biofilm-associated MAC have been shown to have lower susceptibility to antibiotics *in vitro^7^*. For some clinical isolates of *Mycobacterium avium*, virulence seems to be linked to their ability to form biofilms^2^. Compared with isolates of many different pathogenic NTMs, MAC formed the most biofilm in low-resource *in vitro* tests^18^ surpassing *Mycobacterium abscessus*.

In *in vitro* experiments, MAC biofilms have been linked to premature macrophage apoptosis^19^ and better invasion of MAC through airway epithelial cells^20^. In the absence of biofilms, macrophages are known to phagocytose MAC^21^, but phagocytosis is also affected by biofilms. Mature biofilms can prevent phagocytosis^19^, but phenotypic precursors to biofilms, known as microaggregates, have been shown to increase phagocytosis^22^.

After phagocytosis, bacteria can reside and replicate intracellularly as macrophages attempt to kill them^21^. Despite evidence, the role of MAC biofilm and its interaction with bacteria and host immune cells in establishing infections remains unclear^23^. The goal of this current work is to examine the interactions between the innate immune system, bacterial phenotypes, and biofilms (**Figure 1**) in an airway environment. Through this systems analysis we aim to better understand the events in the first 14-days post-inhalation that lead from airway colonization to infection.

**Figure 1.**
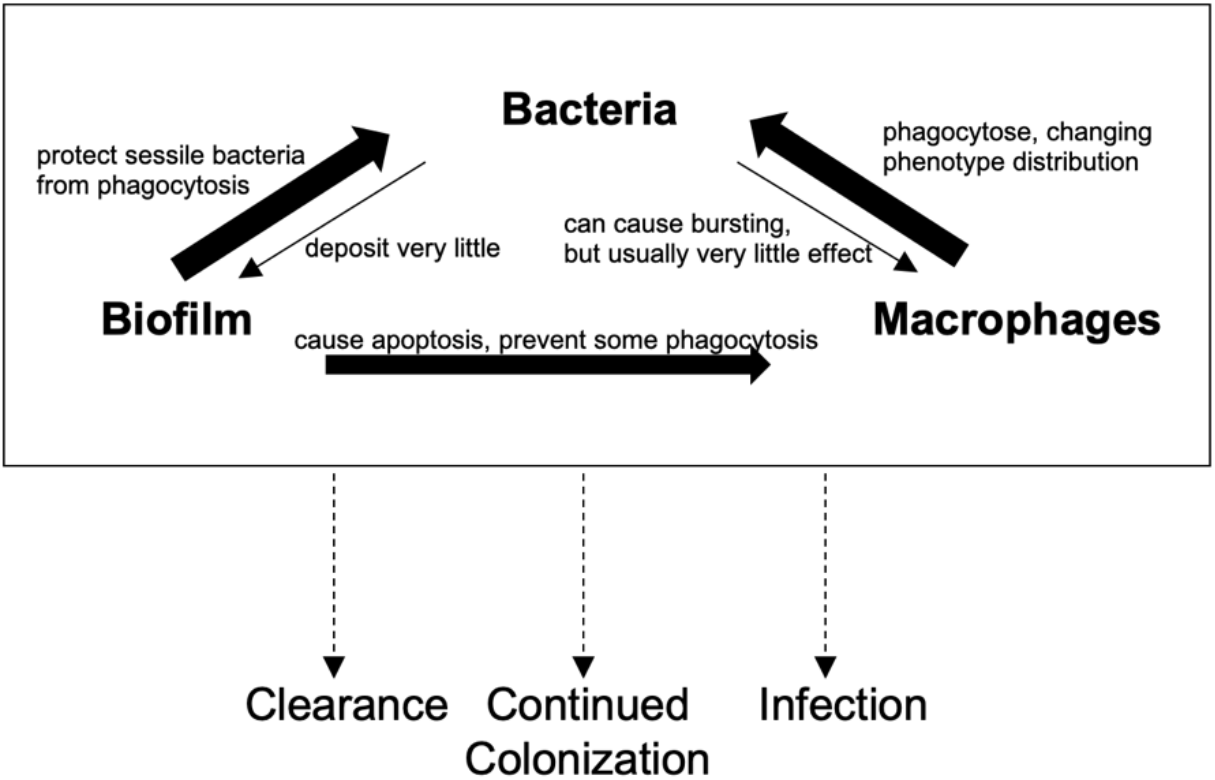
Effects of MAC bacteria, their biofilms, and host macrophages on each other, with effects weighted by arrow size. These dynamics lead to either bacterial clearance, continued colonization of the lung airway, or MAC infection as bacteria move further into the lung.

MAC pulmonary disease typically manifests as nodular bronchiectatic or cavitary disease. NTM-PD is classically described in nonsmoking, post-menopausal female patients^24^. In nodular bronchiectatic NTM-PD, about half of patients have spontaneous sputum conversion^25^, and 88% of patients experience favorable antibiotic treatment outcomes^24^. In contrast, cavitary granulomas always require antibiotic treatment^26,27^. For patients with cavitary disease, favorable outcomes were only observed for 76-78% of patients^24^. Despite common risk factors between patient groups, the mechanisms leading to one type of infection over another remains unclear. Studies with *Mycobacterium tuberculosis* suggest that early interactions with innate immune cells can drive infection progression^28–30^, and we hypothesize that NTM-PD progression to either nodular bronchiectatic or cavitary disease is determined by early interactions between bacteria, biofilm, and innate immune cells in the airways.

These early events in infections are difficult to study in patient populations due to the absence of clinical signs early in disease course, and invasiveness of testing required to study airway colonization. These infections are also poorly replicated in animal models, which often require intravenous injection to establish infection^31,32^. In murine models successfully infected via aerosol, mice often developed disseminated infections with bacteria commonly traveling to the spleen^33^, rather than remaining in the lungs as seen in humans. These challenges lead to the majority studies in patients being observational, reporting the outcomes of treatments with little information on the early events after bacteria are inhaled and deposited in the lungs leading to infection. With challenges of studying these infections *in vivo* and difficulty in replicating the complex environment *in vitro*, computational models are promising tools to obtain a better understanding of early infection events that lead to NTM-PD.

Agent-based models (ABMs) are spatial and temporal stochastic models in which individual “agents,” host immune cells and bacteria, each with unique actions and characteristics interact in an environment by following a specific set of rules^34^. These rules are determined by biological data from experiments that isolate individual interactions (e.g. apoptosis) as well as our understanding of tissue level outcomes (e.g. infection progression and epithelial invasion). Though each rule is relatively simple, the rules combine to form a network of interactions, allowing even simple models to elucidate complex population-level emergent behaviors^34^. Because of their spatio-temporal organization and combinations of simple rules, ABMs are well suited for the examination of interactions between bacteria, biofilms, and immune cells. Such computational models have been used with success in the field of tuberculosis^35^, wound healing^36^, and bacterial biofilm dynamics^37^. Here we describe our new ABM populated with bacteria and macrophages that follow rules determined by *in vitro* experiments. We apply our model to characterize early dynamics of NTM biofilms, lung tissue invasion, and host response in the airway. A better understanding of these early events could elucidate causes of infections and frequent re-infections and may lead to a better understanding of the role of biofilms and novel preventative measures.

## 2. Model description

We built our model in Java using Repast Simphony 2.5.0^38^, and developed two main agent classes: immune cells and MAC bacteria. Agents exist in a discretized 3-dimensional grid, where they interact with each other and their environment over time. Immune cell agents include macrophages in various states: healthy, infected, and apoptosed (dead but not yet cleared from the environment). MAC agents include subclasses of planktonic, sessile, and intracellular bacteria. Time is discretized into “ticks” that represent six-minute time intervals, in which each agent performs its behaviors in randomized order. Interactions between the agents and the environment are described below (parameters are italicized).

### 2.1 Simulation Environment

The spatial environment represents a section of lung airway with mucus coating the inner layer (**Figure 2a**). With grid dimensions of 100×100×3 grid compartments, this mucus is the space in which all agents interact, and the z=0 layer is considered to be the epithelial cells. Each grid compartment represents a space 20×20×20μms for a total space dimension of 2mm×2mm×60μm. The grid is single occupancy for immune cells but multi-occupancy for bacteria.

**Figure 2.**
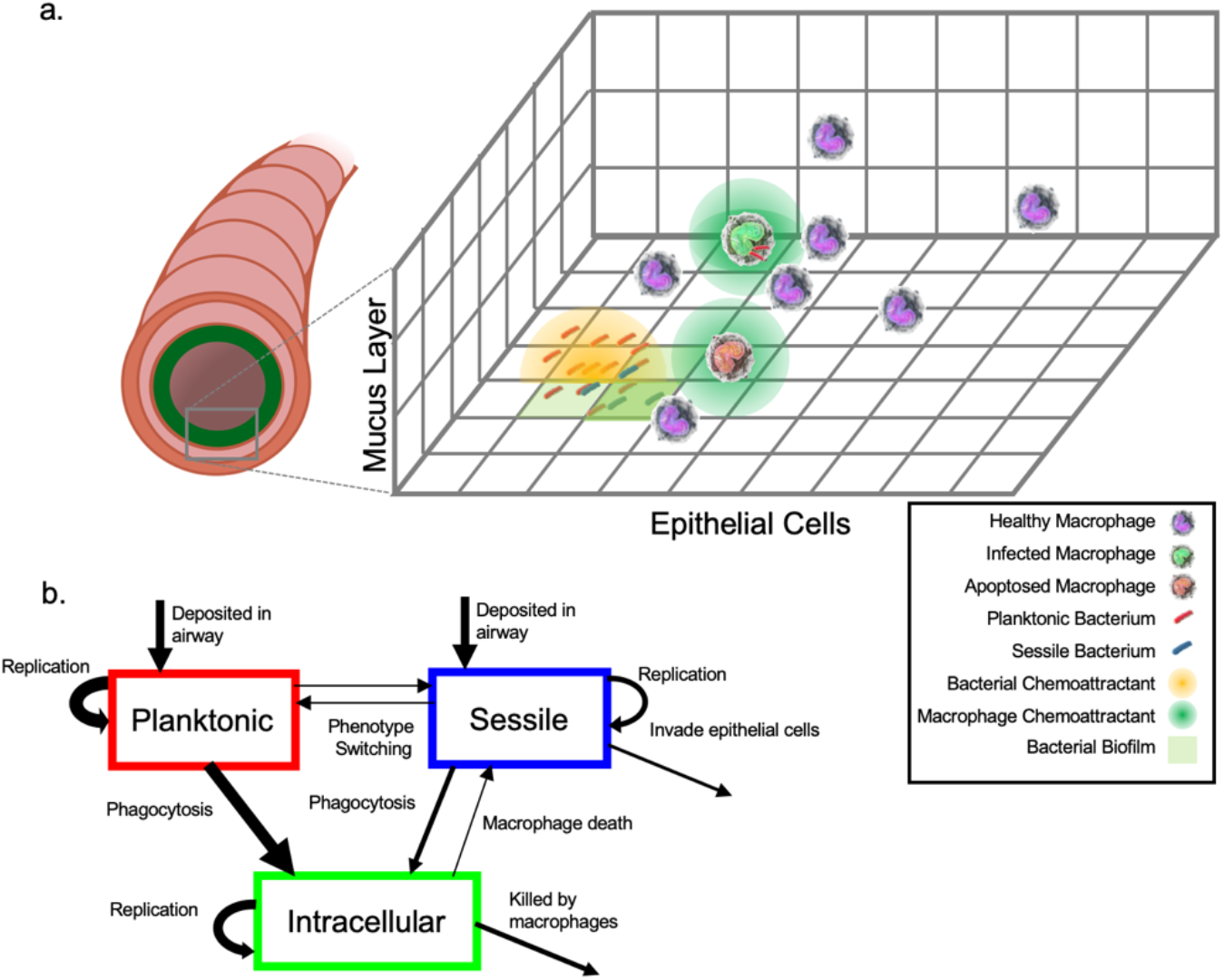
Simulation Layout. a. The grid in which agents operate represents a flattened section of lung airway, with epithelial cells lining the bottom of the grid. Bacteria are randomly deposited in the airway. Sessile bacteria are deposited with some biofilm. Both planktonic and sessile bacteria release bacterial chemoattractant, which causes macrophages to undergo chemotaxis towards them. Once macrophages have bacteria in their Moore neighborhood, they can probabilistically phagocytose them, becoming infected. They may also apoptose in response to biofilms. Both apoptosed macrophages and infected macrophages release macrophage chemoattractants, probabilistically recruiting both patrolling macrophages and additional macrophages from the vasculature. b shows the dynamics of planktonic, sessile and intracellular bacterial phenotypes and the methods by which and individual bacteria can switch between phenotypes, with arrows scaled by probability of occurrence.

The ubiquity of MAC in the environment indicates that most people are exposed, but a significantly smaller fraction of the population actually becomes infected. Instead, these opportunistic pathogens tend to infect patients with pre-existing conditions that impair lung mucosal clearance. For instance, bronchiectasis and chronic obstructive pulmonary disease (COPD) each increase patient risk of contracting NTM pulmonary disease by 187.5 or 15.7-fold, respectively^33^, indicating that mucosal clearance is important in normal immune response to mycobacterium. In this work, we assume that the mucus in our model is nonmoving, therefore representing these comorbid conditions.

We simulate early infection events up to fourteen days post exposure. The adaptive immune system typically does not react until sufficient cytokine signaling from macrophages^39^. In *Mycobacterium tuberculosis* infections, this response has been shown to be further delayed in comparison to other bacterial infections, taking weeks to fully mount a response^40^. With this information, we assume some delay in adaptive immune response to MAC as well, and therefore exclude adaptive immune responses in this version of the model. We assume that this section of airway is similar to adjacent sections and therefore use toroidal boundaries in the x-y plane, and no-flux boundaries in the z direction.

### 2.2 Bacteria

MAC bacteria and their biofilms have been observed in soil, dust, and lakes^5,41,42^, and in man-made environments, such as pipes^43^, showerheads^5^, water filters^44^, pools and hot tubs^45^, and in hospital settings such as catheters^7^ and heater-coolers^8^. Most infections likely occur when patients inhale aerosolized bacteria from common water sources. Mycobacteria have been shown to exist in a 1,000-fold increase of cell density in aerosolized drops from the original water source and to be in particles of a respirable size^46^.

Inhaled bacteria are deposited in the airways. These bacteria may be either planktonic bacteria that have been released into the environment, or sessile bacteria in biofilms that were sheared from pipe walls by water movement, as demonstrated in other biofilm-forming bacterial colonies^47^. Because there are multiple potential inoculum sizes and phenotypes, we simulate colonization for a variety of dose sizes of inhaled bacteria in the lungs. We also test a wide range of amounts of biofilm deposited with each sessile bacterium sheared from the pipe wall.

We initiate a simulation by distributing bacteria uniformly randomly within the center 80% of compartments of the grid where they begin to grow and replicate. Initial bacterial agents have two different extracellular phenotypes: planktonic or sessile, discussed below. The total number of bacteria initially added and the fraction of planktonic versus sessile bacteria initially added is determined by the parameters *N_itlBac*, and *F_itlPlank*, respectively. The base growth rate for any given phenotype is calculated

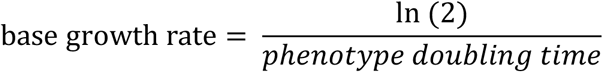

as where phenotype doubling time is given by *T_plankBacDouble*, *T_sessBacDouble*, or *T_ICBacDouble* for planktonic, sessile and intracellular bacteria, respectively. A variance percentage (*S_bacGrowth*) is added or subtracted from this baseline value to ensure diversity amongst the bacterial population.

Though grid compartments are multioccupancy for bacteria, there is a carrying capacity (*K_bacPerGridsquare*) determining how many can occupy a single space that is enforced through logistic growth rates. For each bacterium in each six-minute time step, that bacterium’s biomass is updated as follows:

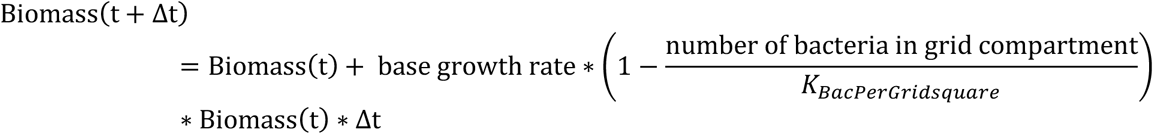

Once a bacterium doubles its biomass, it divides, creating a new bacterium of the same phenotype. After dividing, the biomass of the parent bacteria is reduced by one-half, plus or minus a randomly sampled fraction between 0 and *S_bacDiv*, to add variance and spread out the time to replication for the resulting bacteria^48^. Extracellular bacteria that divide are spatially offset in the z-dimension by a randomly sampled value between negative one and one times the *M_bacZOffset*, and a random angle and some radius between one and three times the *M_bacXYRadius* for the x- and y-dimensions.

Though some research has been conducted on MAC phenotypes and their effects on antibiotic susceptibility and invasion of epithelial cells via biofilm, there is little information on their phenotypic roles in infection. Many types of bacteria take advantage of diverse phenotypes to serve different roles and protect the colony and to communicate via quorum sensing. In this model, we assume that only growth rates and sessile bacteria’s deposition within and ability to make a biofilm are different between the two different phenotypes. The bacterial dynamics, and movement between phenotypes is summarized in **Figure 2b**.

#### 2.2.1 Planktonic Bacteria

Possible behaviors of planktonic bacteria include growing, dividing, and releasing chemoattractant (discussed further in section 3.3). Planktonic bacteria also have some probability of switching phenotypes (*P_phenoSwitch*) each tick if there are any planktonic bacteria in their grid compartment.

#### 2.2.2 Sessile Bacteria

Sessile bacteria are initialized with some amount of biofilm, given by *A_itlBiofilm*, on the assumption that they were sheared from biofilm before inhalation. Sessile bacteria also contribute to the chemoattractant layer, grow, and divide, albeit at a slower rate compared to planktonic and intracellular bacteria. Similar to planktonic bacteria, sessile bacteria can switch phenotypes, depending on *P_phenoSwitch* and if there are sessile bacteria in their grid compartment. If sessile bacteria switch phenotype, they are no longer considered ‘in’ the biofilm in the model and grow more quickly.

*In vitro* experiments have shown that microaggregates, the precursors to sessile bacteria in biofilms, can induce phenotype switching, increase early biofilm production, and are more prone to be phagocytosed^22^. The distinction between microaggregate and sessile phenotypes has not been fully characterized, however, so for the sake of simplicity we consider only one sessile phenotype.

#### 2.2.3 Biofilm

We represent biofilm as a continuous non-diffusing spatial variable in the model. Each grid compartment contains a biofilm value between 0 and 100 arbitrary units (0 representing no biofilm, and 100 representing mature biofilm). Sessile bacteria each contribute a fixed amount of biofilm (*A_bacToBiofilm*) to their grid compartment each tick.

#### 2.2.4 Epithelial invasion

In an *in vitro* study using human bronchial epithelial cells, biofilm-inhibited mutants of a lab strain of MAC were shown to invade at significantly lower rates than their biofilm-forming counterparts and had decreased ability to cause infection in mice in *in vivo* experiments^20^. Therefore, the invasive phenotype is modelled as a boolean variable for all bacteria, with the probability *P_bacInvPheno* to the bacterium when it is created. Only sessile bacteria in biofilm with an invasive phenotype are able to invade. Probability of invasion at a given time is a function of the amount of time that bacterium has been in biofilm (*P_sessInv* multiplied by the number of ticks that bacteria has been in biofilm). In this model version, once a bacterium invades the epithelium it is counted and removed from the environment.

#### 2.2.5 Intracellular Bacteria

Intracellular bacteria are those that have been phagocytosed, but not killed, by a macrophage. Within the macrophage, intracellular bacteria continue to grow and divide. When that infected macrophage bursts or undergoes apoptosis, each bacterium’s probability of being released as a sessile bacterium is determined by *P_bacSurvApop*. These bacteria that are released from inside cells maintain a sessile phenotype due to their reduced growth rate and increased apparent robustness to macrophage killing resulting from intracellular stresses^49^.

### 2.3 Continuous Variables

Continuous variables, where each grid compartment has some value, are used to describe environmental factors and their diffusion within the mucus in the lung airway. These factors include biofilm, bacteria chemoattractant (representing PAMPs) and macrophage chemokines. These values can be detected and altered by the agents.

Though biofilm does not diffuse through the environment, both the bacterial chemoattractant and the macrophage chemokines diffuse with coefficients, *D_bacC* and *D_macC* respectively. Diffusion is modelled using a three-dimensional version of the alternating-direction explicit numerical method, with each grid compartment used as a space step^50^. Because this method is only stable if either time or space steps are small enough, this diffusion calculation occurs 1-4 times per tick (depending on the diffusion coefficient), for each diffusible layer between turns for agents^50^. There is also some degradation of the chemokines and chemoattractants, which is calculated by removing a percentage, *C_macDeg* or *C_bacDeg*, for each diffusion step. Each tick, each extracellular bacterium adds an amount (*A_bacToC*,) to the bacterial chemoattractant value in its grid compartment, and each infected macrophage adds an amount (*A_infMToC*) to the macrophage chemokine value in its grid compartment.

### 2.4 Macrophages

Our macrophage agents represent the innate immune response during the initial colonization and infection of the airway. To begin the simulation, *N_itlMac* macrophages are initialized at random locations throughout the environment, which is based on the density of intra-alveolar macrophages found in the airway in nonsmokers^51^. Based on interactions with other agents and the environment, macrophages are further classified as healthy, infected, and dead, which affect the cell’s behaviors.

One notable role of biofilms deposited in the environment by sessile bacteria is the apoptotic effects they have on macrophages^19^. Rose and Bermudez showed that, when healthy phagocytic THP-1 cells are placed on fully-formed avium biofilms that a large number apoptose, many within 30 minutes of exposure. In biofilms both with and without live *Mycobacterium avium* bacteria, macrophages produce high levels of signaling molecules TNF-α and nitric oxide, leading to high levels of apoptosis without a reduction to either biofilm volume or bacteria population^19^. *In vivo*, this signaling would lead to high levels of inflammation and impaired lung function with little-to-no bacterial killing.

We simulate this macrophage sensitivity to biofilms by assigning each macrophage an apoptotic signal tolerance. Each macrophage will apoptose when it is exposed to biofilm levels above its tolerance. The distribution of macrophage tolerance is based on the *in vitro* data discussed above^19^ (**Supplement 1**). When the macrophage undergoes apoptosis it kills a fraction of the internal bacteria (each bacterium has probability, *P_bacSurvApop*, of surviving) and leaves behind a dead macrophage. Macrophages can recover some of their biofilm tolerance when they are not exposed to biofilm.

#### 2.4.1 Healthy Macrophages

Alveolar macrophages patrol the lower lung airway mucus. There, they undergo chemotaxis to find and phagocytose foreign particles (including bacteria) that have been deposited in the airway mucus^52^. In our chemotaxis algorithm, the direction of cellular movement is determined by summing the values of both the bacterial chemoattractant and macrophage chemokines for each grid compartment in the macrophage’s Moore neighborhood. We calculate the probability of the macrophage moving to each grid compartment by dividing the chemoattractant concentration for that grid compartment by the total chemoattractant for all grid compartments in the Moore neighborhood (**Supplement 2**). Macrophages can move by one grid compartment per tick, and select their direction randomly according to the probabilities calculated. If the sum of the chemoattractant and chemokine concentrations in the macrophage’s Moore Neighborhood is below the *A_threshChemotax*, the macrophage will move in a random direction.

After moving, macrophages have the ability to phagocytose bacteria located within their Moore neighborhood. Mycobacterial biofilms have been shown to prevent or slow phagocytosis and overstimulate innate immune cells leading to apoptosis. This effect has been shown in *in vitro* experiments, where THP-1 cells were unable to reduce bacterial population levels in pregrown biofilm. The model reflects this phenomenon when macrophages select a bacterium to phagocytose. Macrophages first create a list of bacteria in the Moore neighborhood. They add all planktonic bacteria in their Moore neighborhood to this list, but the probability of each sessile bacterium being added to the list is (100 - biofilm level)% to mimic the protective properties of biofilm. A single bacterium is then randomly selected from the list and has some probability (*P_mPhag*) of being phagocytosed. If phagocytosis is successful, this process is considered an infection event, and the healthy macrophage becomes infected.

#### 2.4.2 Infected Macrophages

Infected Macrophages no longer move, but may continue to phagocytose bacteria that are in their Moore neighborhood, with a reduced probability *P_infMPhag*. Each tick they also have some probability (*P_macKillsICBac*) of killing an internal bacterium.

If an infected macrophage is able to kill all of its internal bacteria, it becomes uninfected, and converts back to being a healthy macrophage. However, if the number of bacteria within the macrophage surpasses the *N_threshBurst*, that macrophage bursts, releasing all bacteria back into the environment. Biologically, this bursting has been hypothesized to help bacteria move around the body in disseminated disease^49^.

New, uninfected macrophages are recruited to the site of infection through bronchial microvasculature^53^, driven by chemokines released by infected macrophages. The cytokines released to induce recruitment commonly include some combination of pro-inflammatory signals, such as tumor necrosis factor-α (TNF-α)^54^ and anti-inflammatory signals including interleukin-10 (IL-10)^55^. In our model, there are a number of “recruitment areas” (*N_recAreas*) throughout the grid which represent blood vessels. Given that the concentration of chemokines released by infected macrophages has reached a threshold at one of these locations (*A_threshRec*), there is some probability (*P_rec*) each tick of a new, healthy macrophage being recruited to that location.

Finally, if continued biofilm exposure causes the infected macrophage to apoptose instead, it kills some of its internal bacteria, and releases a fraction (probabilistically determined by *P_bacSurvApop*) into the extracellular space.

#### 2.4.3 Dead Macrophages

Dead macrophages remain for *T_apopDeg* ticks before being removed from the grid, representing their degradation.

### 2.5 Model Implementation

Parameter ranges are based on literature where available (**Table 1**). While exact dynamics in the airway system remain unclear, we do have physiological boundaries for some parameters (e.g. bacterial doubling times, diffusion rates, macrophage movement). If parameter estimates are not available in the literature initial ranges were estimated by systematically narrowing ranges to eliminate outcomes that are not supported in literature (e.g. predominantly uncontrolled exponential growth within days of infection). We use these biologically feasible parameter ranges to evaluate the distribution of possible infection outcomes. In order to examine the full range of possible behaviors in the model, we use Latin Hypercube Sampling to vary parameters within biologically or physically feasible ranges, then iteratively run simulations to narrow down the ranges, eliminating simulations parameters with instant clearance or no phagocytosis (**Figure 3**). We then re-select parameters, and examine the results of a total of 900 simulations^56^, with 300 parameter sets and three unique replicates for each parameter set. We use partial rank correlation coefficient (PRCC) to analyze uncertainty in various outputs (planktonic, sessile, and intracellular bacteria counts, and healthy and infected macrophage counts) based on uncertainty in different parameter values (α = 0.01).

**Figure 3.**
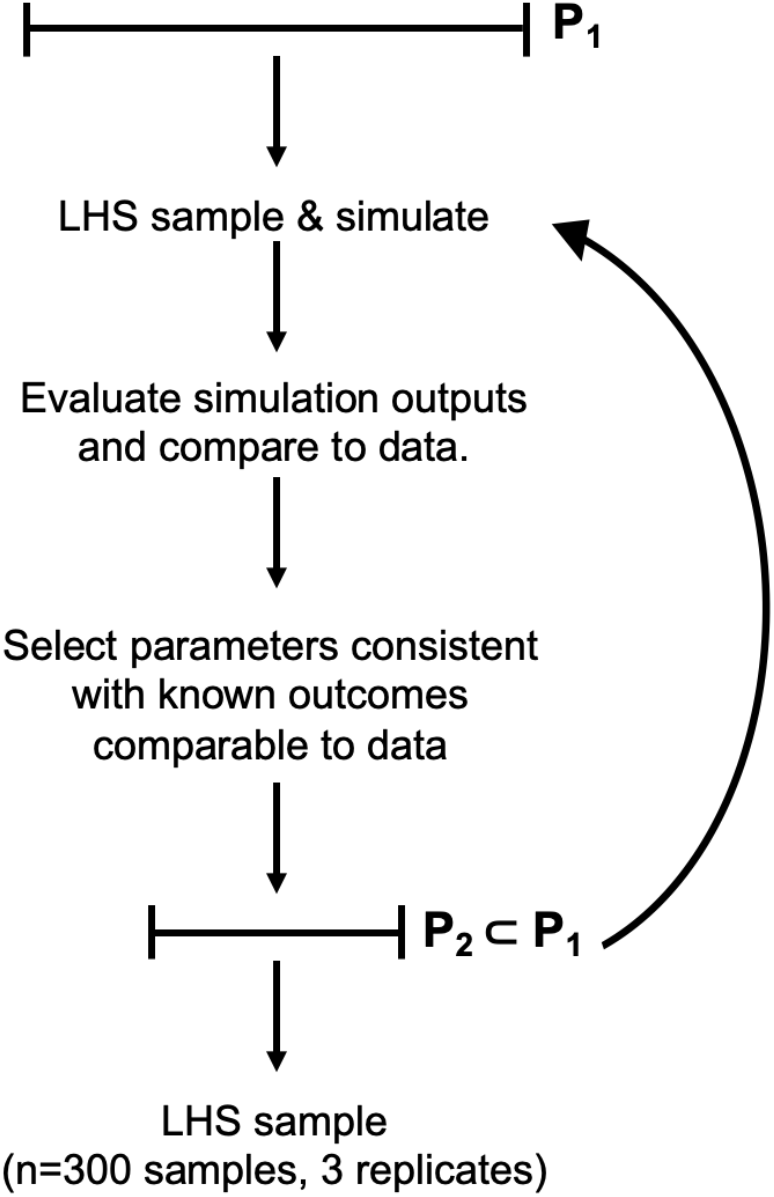
Parameter space was iteratively narrowed to eliminate simulations with immediate bacterial clearance and bacterial growth or macrophage recruitment that is so fast as to be unbiological. Once this narrowing was complete, the final data set was selected via LHS to include 300 samples, and three replicates of each sample with different random seeds, for a total of 900 simulations.

**Table 1.** Parameters were taken from literature when possible. Parameter sources marked with a citation and asterisk indicate further narrowing of the parameter range during calibration, while parameter sources marked “e” indicates that the parameter was estimated, and given a large range due to uncertainty. Finally, diffusion coefficients were independently varied, but were calculated based on the diffusion rate of interleukin 6 (IL-6) in water^59^ divided by a value between 38 and 270 to account for the increased viscosity of mucus in diseases such as COPD and Cystic Fibrosis^60^.

| Parameter Name | Description | Range | Unit | Source |
| --- | --- | --- | --- | --- |
| N_itlBac | Initial number of bacteria deposited | 1 – 50 | Scalar | e |
| F_itlPlank | Fraction of initial bacteria that are planktonic | 0.2 - 0.8 | Fraction | e |
| A_itlBiofilm | Amount of biofilm per initial sessile bacterium | 1 – 100 | Scalar | e |
| N_itlMac | Initial number of macrophages | 60 – 130 | Scalar | <sup>51</sup> |
| N_recAreas | Number of potential recruitment areas | 450-909 | Scalar | <sup>57</sup> |
| T_apopDeg | Time for dead macrophages to be removed/degrade | 0.5 – 72 | Hours | e |
| C_bacDeg | Fraction of total bacterial chemoattractant that degrades each tick | 0.0001 – 0.1 | %/6 min | e |
| D_bacC | Bacterial chemoattractant diffusion coefficient | 1E-9 – 7E-9 | cm<sup>2</sup>/s | e |
| P_bacSurvApop | Probability of an individual IC bacterium surviving in apoptosis | 20 – 70 | % | e |
| P_bacInvPheno | Probability of an individual bacterium having an invasive phenotype | 6 – 15 | % | <sup>20</sup> |
| K_bacPerGridsquare | Bacterial carrying capacity per gridsquare | 20 – 300 | Scalar | e |
| T_ICBacDouble | Doubling time for IC bacteria | 100 – 140 | Hours | <sup>58</sup> |
| A_bacToC | Individual bacterium's contribution to bacterial chemoattractant per tick | 5 – 30 | scalar | e |
| M_bacZOffset | Distance a child bacterium is offset from parent bacteria in z-dimension in extracellular replication | 0.006 – 0.016 | cm | e |
| M_bacXYRadius | Distance child bacteria is offset from parent bacteria in xy-dimension in extracellular replication | 0.0006 – 0.0016 | cm | e |
| T_plankBacDouble | Doubling time for planktonic bacteria | 20 – 39 | Hours | <sup>16*</sup> |
| P_phenoSwitch | Probability of an extracellular bacterium switching phenotypes given a bacterium of the opposite phenotype is in the same gridsquare, per tick | 0.01 – 0.1 | % | e |
| T_sessBacDouble | Doubling time for sessile bacteria | 40 – 80 | Hours | <sup>16*</sup> |
| P_sessInv | Probability of a sessile bacterium in biofilm invading the epithelium | 0.001 – 0.1 | % | <sup>20*</sup> |
| S_bacDiv | Variance in bacterial division | 10 | % | e |
| S_bacGrowth | Variance in bacterial growth rates | 10 | % | e |
| A_bacToBiofilm | individual sessile bacterium's contribution to biofilm per tick | 1E-6 – 1E-3 | Scalar | <sup>19*</sup> |
| P_infMPhag | probability of an infected macrophage phagocytosing a bacterium, per tick | 10 – 50 | % | e |
| A_threshChemotax | Amount of total chemoattractant necessary for a macrophage to chemotax | 0.1 – 10 | Scalar | e |
| A_threshRec | amount of macrophage chemoattractant necessary at recruitment area to activate | 20 - 1000 | scalar | e |
| N_threshBurst | number of IC bacteria necessary to cause an infected macrophage to burst | 10 - 60 | scalar | <sup>58</sup> |
| C_macDeg | Fraction of total macrophage chemoattractant that degrades each tick | 0.0001 – 0.1 | %/6 min | e |
| D_macC | macrophage chemoattractant diffusion coefficient | 1E-9 – 7E-9 | cm<sup>2</sup>/s | e |
| A_infMToC | individual infected macrophage's contribution to macrophage chemoattractant per tick | 30 – 100 | Scalar | e |
| P_macKillsICBac | probability of infected macrophage killing an intracellular bacterium, per tick | 0.01 – 0.1 | % | e |
| P_mPhag | probability of healthy macrophage phagocytosing a bacterium, per tick | 50 – 100 | % | e |
| P_rec | probability per tick of a macrophage being recruited at the recruitment area, given the threshold to recruit has been met | 0.01 – 1 | % | e |
| areaDim | number of gridsquares in the x or y direction | 100 | scalar |  |
| zDim | number of gridsquares in the z direction | 3 | scalar |  |

## 3. Results

### 3.1 Most MAC exposures result in predominantly intracellular bacterial populations

We first aim to characterize the dynamics of bacterial phenotypes following MAC exposure. Our model predictions indicate early and effective phagocytosis of extracellular bacteria. The total bacterial load is stable or increasing for 536 of the 900 simulations (59.6%) within the first 24 hours following exposure. These dynamics are the result of a decrease in both planktonic and sessile bacteria within the first 24 hours, and a corresponding increase in intracellular bacteria (**Figure 4a-c**). In 50.8% (n=457) of simulations, macrophages phagocytose all planktonic bacteria within the first 24 hours, and in 50.9% (n=458) of simulations macrophages phagocytose all sessile bacteria within the first 24 hours. Overall, in 33.8% (n=304) of simulations all extracellular bacteria were phagocytosed in the first 24 hours.

**Figure 4.**
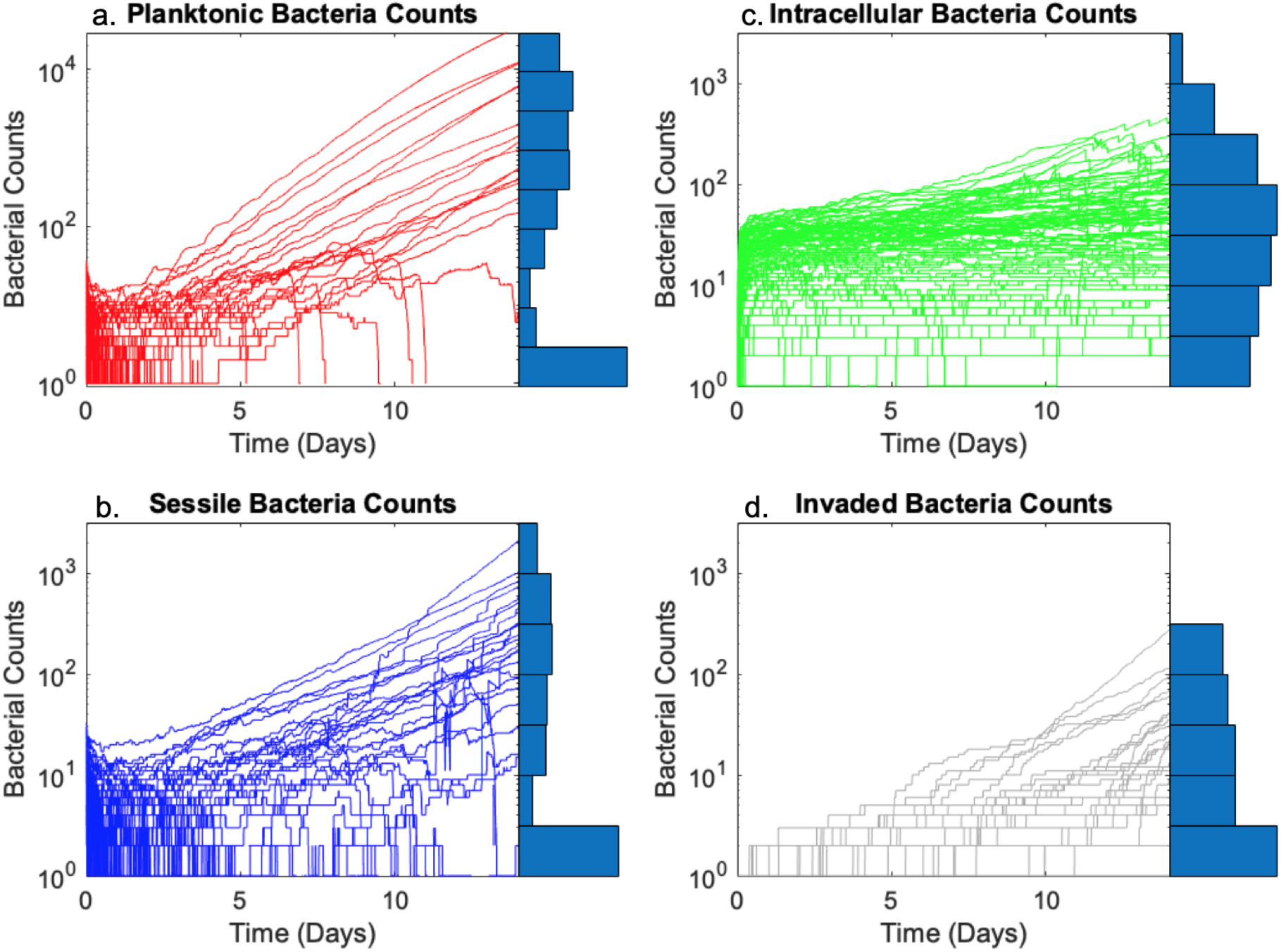
Timeseries plots of the bacterial counts for a) planktonic, b) sessile, c) intracellular, and d) bacteria that invaded the epithelium for 100 randomly selected simulations. The distribution of final counts for all 900 simulations is shown to the right of each graph, with maximum amplitude normalized to one, then converted to logarithmic scale. Note, due to higher bacterial loads, y-axes labels are different for panel a, showing planktonic bacterial counts.

Following the initial phagocytosis events, the number of intracellular bacteria increases more slowly (**Figure 4c**), despite bacterial killing by macrophages, due to continued replication of intracellular bacteria and phagocytosis. In the 66.2% (n=596) of simulations with remaining extracellular bacteria after 24 hours, individual bacteria begin to grow exponentially, but the population can still be phagocytosed, leading to a majority of these simulations also having mostly intracellular bacteria. As the number of sessile bacteria increase, so does the number of bacteria predicted to have invaded through the epithelial layer (**Figure 4d**).

Uncertainty analyses confirm the impact of macrophages effectively finding and phagocytosing bacteria on controlling bacterial phenotypes. The amount of chemoattractant produced by each individual bacterium (*A_bacToC*) and the rate of chemoattractant degradation (*C_bacDeg*) both affect the chemoattractant concentrations and gradients; and both are significantly correlated with bacterial numbers. Higher chemoattractant production (*A_bacToC*) and lower chemoattractant degradation (*C_bacDeg*) enable macrophages to find and phagocytose bacteria more efficiently and therefore initially lead to higher intracellular bacterial numbers and lower extracellular bacterial numbers (day 0 to 4). However, this more efficient phagocytosis leads to lower bacterial numbers both intra- and extra-cellularly after days 4-6 as macrophages are able to kill intracellular bacteria.

In those early timepoints (4 hours), macrophages finding bacteria to phagocytose is largely a matter of chance, and the probability increases higher macrophage densities (discussed in Section 3.2). After 4 hours, macrophages follow the gradients of chemoattractants produced by the bacteria. At this point we observe a significant negative correlation (PRCC) between the amount of chemoattractant individual bacteria (*A_bacToC*) and both planktonic and sessile bacterial counts (**Supplement 6**), and a significant positive correlation between the rate of chemoattractant degradation (*C_bacDeg*) and these bacterial counts (**Supplement 6**). These parameters both control the amount of total attractant and the gradient that determines how “easily” macrophages find the bacteria.

To better understand the balance between bacterial phenotypes within individual simulations, we evaluate each simulation based on the proportion of bacteria that are planktonic, sessile and intracellular over time (**Figure 5**). Overall, 30 (3.3%) simulations clear all bacteria. The remaining simulations have predominantly intracellular bacterial populations for the first 4-8 days, after which three broad categories of infection dynamics emerge. The first and largest category (n=735, 81.7%) includes those simulations with predominantly or exclusively intracellular bacteria (n=81, 9% and n=654, 72.7% of total runs, respectively). Closer investigation of simulations in this predominantly intracellular category show that many macrophages are nearing burst thresholds, indicating that the system is still in flux, which would have implications for infection dynamics after 14 days when adaptive immune responses become active.

**Figure 5.**
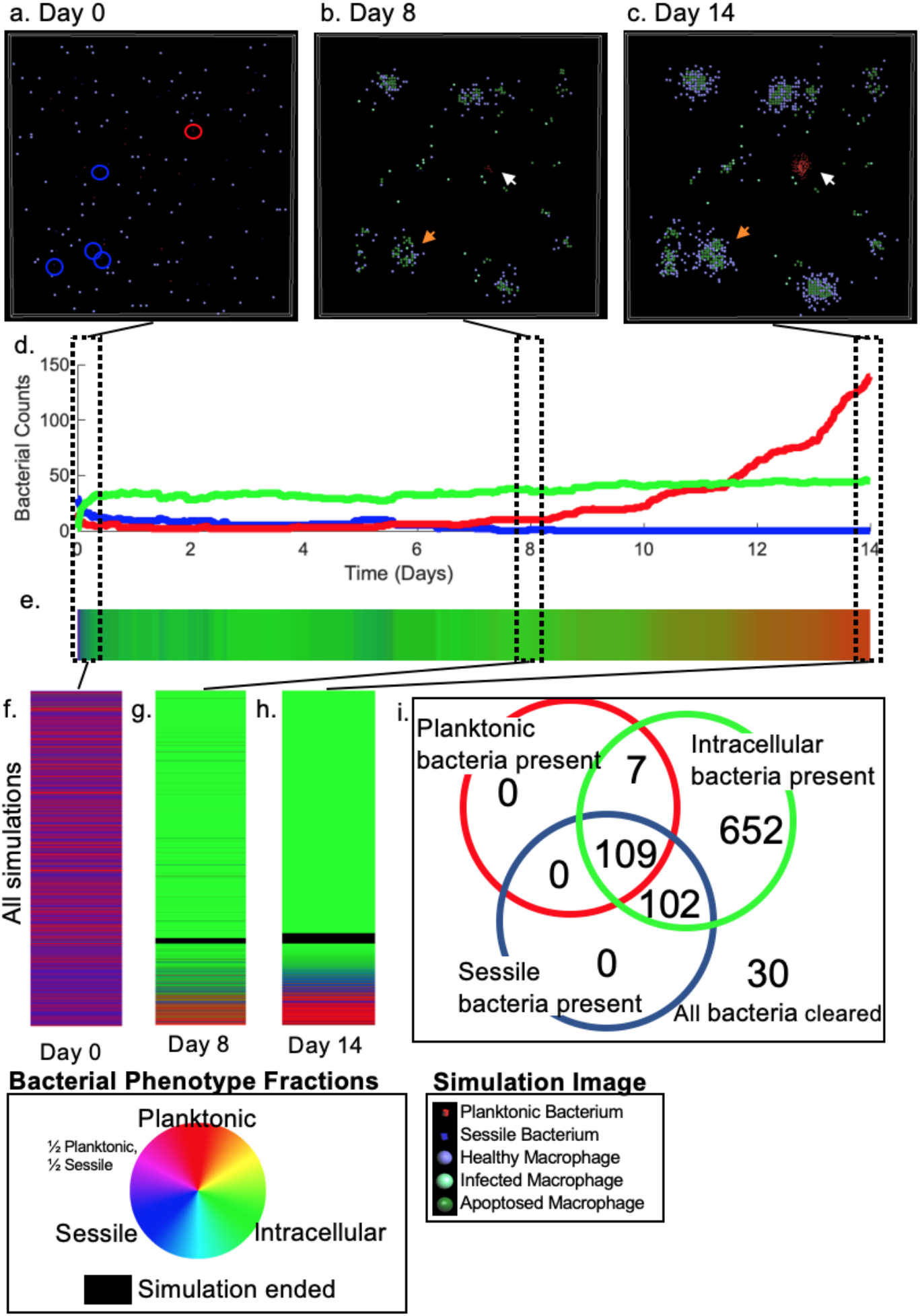
Images from simulations and analyzing phenotype distributions. For one representative simulation, a) shows the initial deposition of bacteria in the airway (a few are circled in red and blue for planktonic and sessile bacteria, respectively), with patrolling healthy macrophages (purple). b) shows day 8, highlighting the effects of spatial distributions of macrophages vs. bacteria. The orange arrow indicates a single infected macrophage (bright green) surrounded by macrophages it recruited, some of which have apoptosed (dark green) due to biofilm exposure. The white arrow shows a cluster of planktonic bacteria that have remained extracellular, and are replicating quickly. c) shows the end of the simulation (day 14). The orange arrow shows further recruitment and apoptosis of macrophages around an infected macrophage. The white arrow shows the planktonic bacteria colony that has grown. For the same representative simulation, d) shows the bacterial counts for planktonic (red), sessile (blue) and intracellular (green) bacteria. In e), the relative proportion of the bacterial phenotypes are plotted in color combinations of red, blue, and green, representing planktonic, sessile, and intracellular bacteria, at each timestep. The color wheel demonstrates how these colors combined to represent the relative proportions. In f-h), we examine the relative bacterial proportions across all 900 simulations at days 0, 8, and 14, where each horizontal line shows the distribution for one simulation at that timepoint. The black line shows simulations that ended after all bacteria were killed. Finally, i) shows the distributions of which bacterial phenotypes are present at the end of the 14-day simulation, regardless of proportions.

The second category (n=97, 10.8%) of simulations contains a those with a majority of planktonic bacteria. Here, a few groups of rapidly-growing and spatially isolated planktonic bacteria typically dominate towards the end of the simulation (**Figure 5b-c, white arrow**). These colonies are mostly seeded by a few planktonic bacteria that remain extracellular throughout the entire simulation (**Supplement 3**). In some cases, planktonic bacterial populations are bolstered by sessile bacteria changing to a planktonic phenotype, but this accounts for only 20.1% of the planktonic bacterial population even in the simulation with the most phenotype switching (the median across all simulations is 0.35%). Planktonic bacteria have also been observed to indirectly protect nearby sessile bacteria from phagocytosis, as macrophages prioritize phagocytosis of planktonic bacteria over sessile bacteria. Remaining sessile bacteria are then better able to invade the epithelial layer.

The third and smallest category (n=38, 4.2%) is comprises simulations with a majority sessile bacteria (**Figure 5h**). These populations usually have high levels of biofilm, which protect the extracellular sessile bacteria from phagocytosis. Total biofilm levels in the simulation grid increase linearly in all simulations (**Supplement 4**). However, average local biofilm levels decrease because sessile bacteria spread out faster than they can produce biofilm. Predominantly sessile simulations also have the highest invasion rates, as the number of sessile bacteria at the end of the simulation and total bacteria that invade are significantly linearly correlated (p<.001, r^2^=0.938).

Beyond the predominant phenotypes, the model predicts that certain combinations of phenotypes are more likely than others. Of the 870 simulations that did not clear all bacteria, 218 (24.2%) have more than one bacterial phenotype present at the end of the simulation (**Figure 5i**). Every simulation with remaining bacteria, however, has at least some intracellular bacteria, indicating that the innate immune response continues to play a role in bacterial phenotype distributions despite large extracellular bacterial populations.

Thus, there are several possible outcomes following MAC exposure. In most cases, however, macrophages efficiently phagocytose bacteria yielding mostly intracellular bacteria at the end of 14 days. High bacterial loads are mostly associated with planktonic bacteria. Given the predominance of the intracellular bacterial populations, we next examine macrophage dynamics and their interactions with MAC following exposure.

### 3.2 Sustaining macrophage populations is key to control extracellular bacterial dynamics

Total numbers of healthy macrophages broadly exhibit two trajectories, with 31.6% net increasing (n=284) and 68.2% decreasing (n=614) over 14 days (**Figure 6a-b**). As with intracellular bacteria, we see an early increase in the number of infected macrophages in the first day of the simulation. Unlike intracellular bacteria, the infected macrophage numbers remain relatively stable or decrease as the simulation progresses.

**Figure 6.**
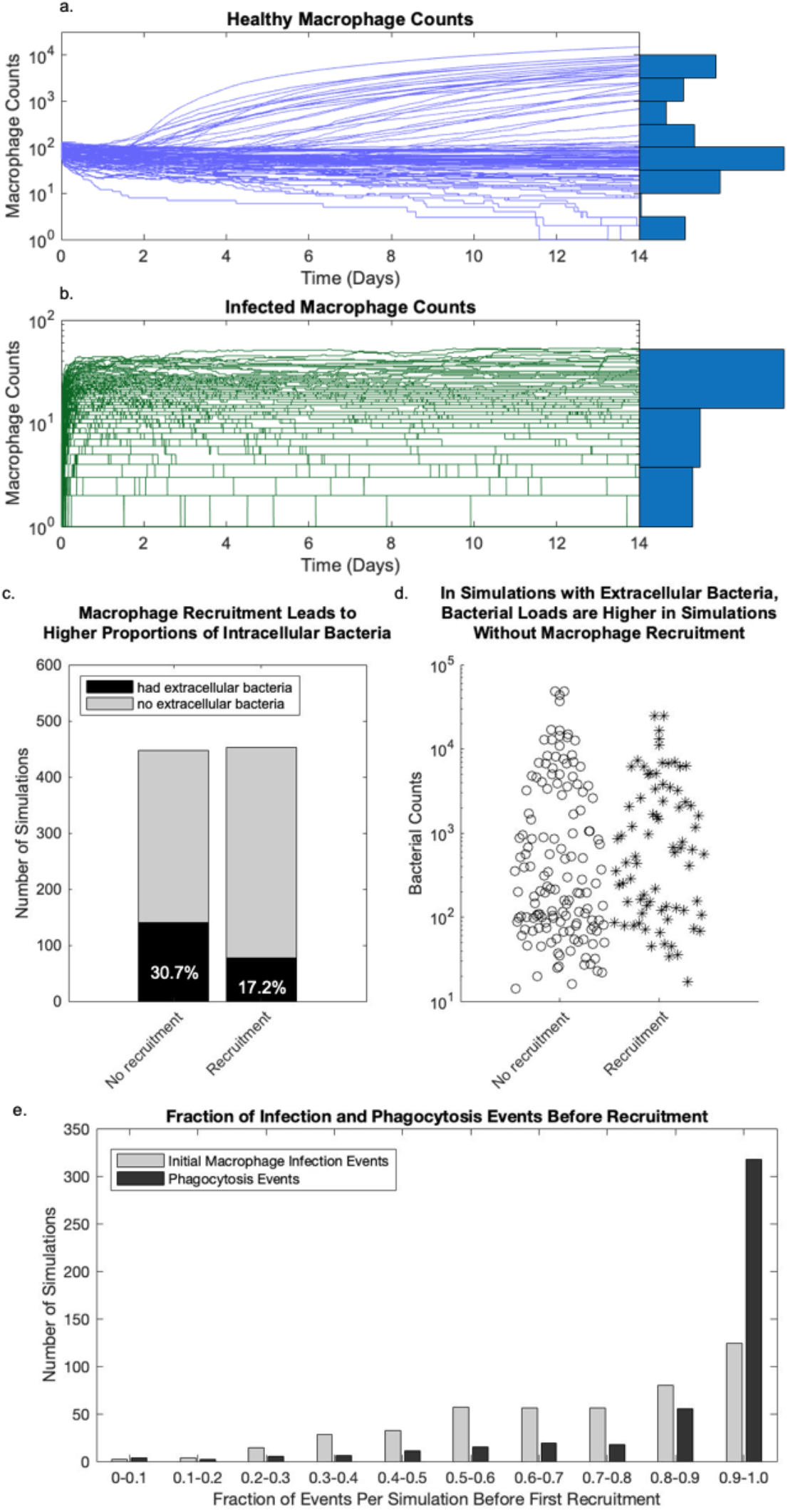
Macrophage dynamics. a-b) show the healthy and infected macrophage count dynamics, respectively, with the distribution of final counts for all 900 simulations on the right of each graph, with maximum amplitude normalized to one, then converted to logarithmic scale. In c) we show the proportion of simulations with remaining extracellular bacteria in simulations with without recruitment. Of those simulations that did have extracellular bacteria, d) shows the distribution of bacterial loads between simulations with and without recruitment. e) shows a histogram of the proportion of both macrophage infection events (the first time a macrophage is infected) and phagocytosis events (every time a bacterium is phagocytosed) before the first macrophage has been successfully recruited.

This sustained population of infected macrophages results from a balance between macrophages becoming infected, and infected macrophage loss due to a combination of macrophage death and infected macrophages uninfected as they kill their intracellular bacteria. Overall, we find that apoptosis of uninfected macrophages due to biofilm exposure accounted for over 50% of macrophage deaths in 695 (77.2%) of the simulations. This trend is also skewed heavily toward even higher percentages, and 90.8% of all macrophage deaths across all 900 simulations are due to apoptosis. This finding suggests that biofilm-induced apoptosis has significant effects on the number of macrophages in the airways in patients with biofilm-forming MAC colonies.

New macrophages are recruited in 49.7% of simulations (447 simulations). In the remaining 453 (50.3% of total) cases there is a failure to recruit more macrophages. Simulations without macrophage recruitment are significantly (p< 0.0001) less likely to eliminate extracellular bacteria (**Figure 6c**); of 453 simulations without recruitment, 139 (30.7%) have remaining extracellular bacteria, compared to the 447 simulations with recruitment where only 77 (17.2%) have remaining extracellular bacteria. For simulations that end with extracellular bacteria, those without recruitment also have higher bacterial numbers compared to those with successful recruitment (**Figure 6d**). Simulations without macrophage recruitment show clusters of extracellular bacteria that are not efficiently contained by those macrophages within the simulation. In simulations with high macrophage recruitment (**Figure 4a-c**), macrophages quickly move towards bacteria and phagocytose many, but not necessarily all, extracellular bacteria. This observation indicates that macrophage recruitment played a large role in suppressing extracellular bacteria populations in our simulations.

However, recruitment was not always sufficient to sustain the macrophage population. In 141 out of 447 simulations with recruitment (31.5%), the combined number of healthy macrophages and infected macrophages still undergoes a net decrease, due to macrophage death. These deaths are especially prevalent in simulations with high numbers of sessile bacteria, due to the pro-apoptotic effects of biofilm. In simulations with a majority of sessile bacteria at the end of the simulation (39 simulations), 38 (97.4%) show overall decreases in total macrophage counts from the beginning to the end of the simulation, even with 10 of those 38 simulations having macrophage recruitment.

Our simulations therefore suggest that macrophage numbers fluctuate due to a combination of recruitment and cell death. While recruitment of macrophages has a negative correlation with remaining extracellular bacteria, recruitment is not necessary in all cases for macrophages to phagocytose all the bacteria, nor a guarantee that no extracellular bacteria will remain. To further explore this connection between macrophage recruitment and bacterial levels we next quantify the impact of macrophage recruitment and movement on bacterial sub-populations.

### 3.3 Despite recruited macrophage counts having a significant negative correlation with extracellular bacteria counts, most recruited macrophages never phagocytose a bacterium

Comparing macrophage recruitment with bacterial dynamics, we find that total recruited macrophages have a significant negative correlation with both day 14 planktonic bacteria (p=0.0108, r^2^=0.00611) and sessile bacteria (p=0.00152, r^2^=0.01). In contrast, total recruited macrophages do not correlate with intracellular bacteria counts (p=0.4). Taken together, the insignificant p-value for intracellular bacteria and the low r^2^ values for planktonic and sessile bacteria indicate that macrophage recruitment does not account for much of the observed variability in the bacterial subpopulations.

The reason macrophage recruitment does not greatly influence bacteria is that recruited macrophages are useful to control bacterial load but only below a certain level of macrophage recruitment. Any additional macrophage recruitment past that point does not reduce the number of bacteria any further. In fact, the majority of simulations show high proportions of initial infection and phagocytosis events occur prior to macrophage recruitment (**Figure 6e**). This timing is somewhat expected, as infection events are necessary to recruit macrophages, but does mean that most recruited macrophages never phagocytose a bacterium.

This failure of newly recruited macrophages to phagocytose any bacteria is due to one of two factors in our simulations– 1) either there are no remaining extracellular bacteria when they are recruited, or 2) recruited macrophages fail to find and phagocytose extracellular bacteria. Of the 447 simulations with recruitment, 370 (82.8%) have no extracellular bacteria by the end of the simulation. Further, this complete phagocytosis is skewed early in the simulation, with this total phagocytosis occurring in the first two days (**Supplement 5**), meaning that there are no extracellular bacteria when many of these macrophages are recruited. In the remaining 77 simulations with remaining extracellular bacteria despite recruitment, 70 (90.9%) of these contain some extracellular bacteria at all times, indicating that extracellular populations are more often sustained by persistent extracellular bacteria, rather than originating from bacteria that are released from infected macrophages. There are also situations in which macrophages cannot access colonies of bacteria, as infected and apoptosed macrophages have formed a barrier around them, preventing access (**Figure 5b-c, orange arrows**).

These results indicate that early phagocytosis of bacteria by resident macrophages, more than continued recruitment of macrophages, have long-term impacts on intracellular bacteria. This finding suggests that the number of macrophages present at the time of exposure is key to keeping bacterial numbers low. We next seek to identify other parameters that impact bacterial numbers longer term.

### 3.4 Initial parameters have large, lasting impacts on infection progression

To examine the factors that affect infection outcomes, such as macrophage and bacterial phenotype numbers over time, we calculate Partial Rank Correlation Coefficients (PRCCs) between each model parameter and each model output over time (**Supplement 6**). We find that initial parameters continue to have a significant impact throughout much of the simulation (**Figure 7a-c,e**).

**Figure 7.**
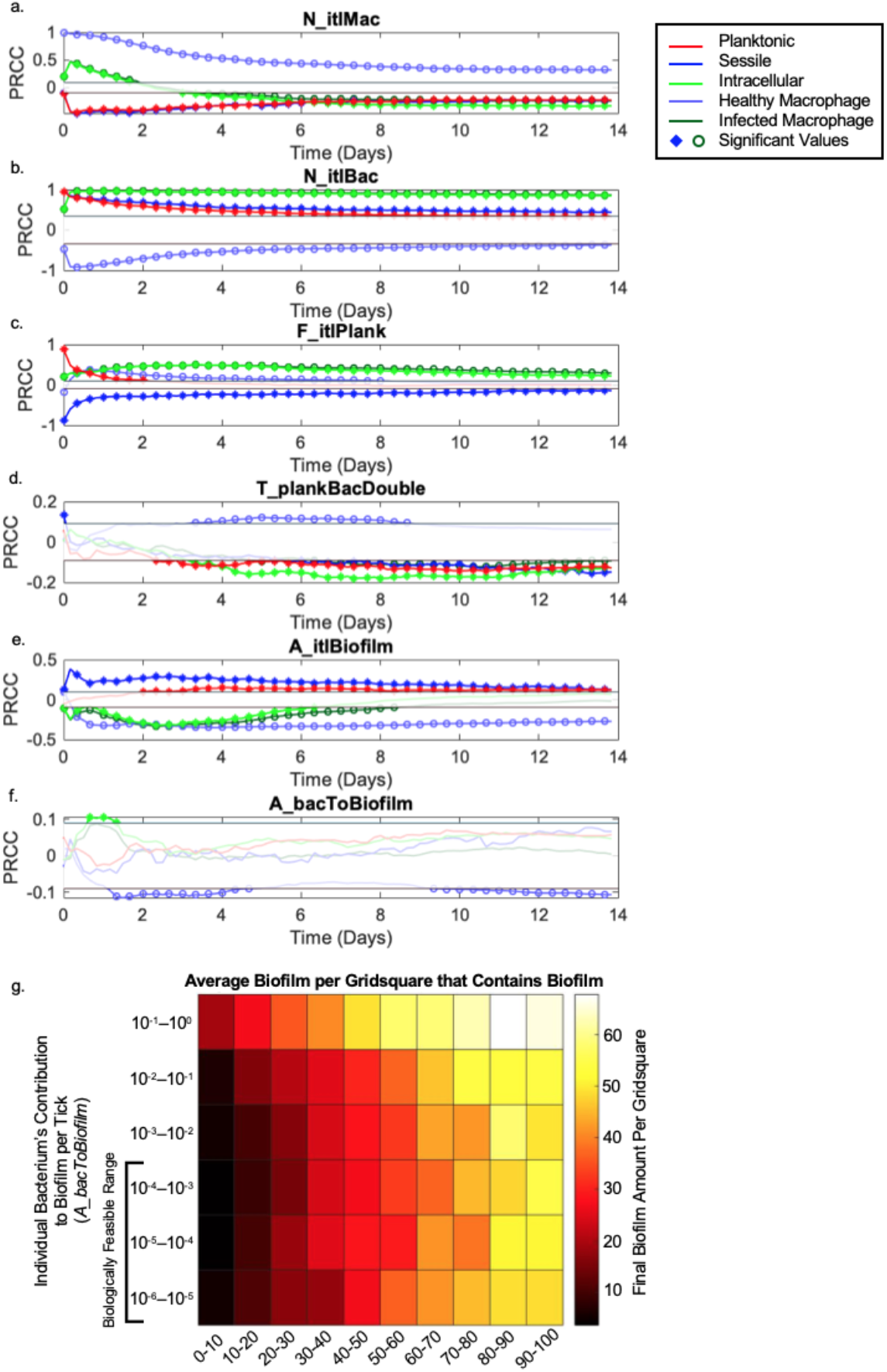
Initial parameters’ effects on model outcomes. a-f show the Partial Rank Correlation Coefficient of each parameter on planktonic, sessile, and intracellular bacterial counts, and healthy and infected macrophage counts. The range in which these values are not significant centers around 0, and the threshold is marked by a horizontal black line above and below. Values in the insignificant range are plotted as lines, but do not have individual data markers and are greyed out. Significant values are marked, using a filled point for bacteria or an open point for macrophages. In each plot, intracellular and infected macrophage coefficients are very similar, and plots often overlap. a) shows the PRCCs for initial macrophage counts, b) the initial bacterial load deposited in the airway, and c) the fraction of those initial bacteria that are planktonic (rather than sessile). d) shows the PRCC values for the planktonic bacteria doubling time (which is negatively correlated with growth rate) for contrast, as a non-initial parameter. e) and f) show the PRCC values for initial amount of biofilm that sessile bacteria are deposited with and the amount of biofilm created by a sessile bacteria per tick, respectively. Finally, in g), we further compare deposited vs. created biofilm. Here, we compare the the average amount of biofilm per gridsquare that contains biofilm at the end of the simulation, which ranges from 0-100%, across initial and contributed biofilm amounts. We used our initial data set, for the “biologically feasible” contributed biofilm range, and an additional data set in which contributed biofilm was increased logarithmically. We see that average biofilm loads are driven largely by initial values, as individual sessile bacteria grow biofilms so slowly that their contributions do not have a significant effect on the average until the contribution is 100x the biologically feasible range.

The initial number of macrophages (*N_itlMac*) is initially strongly, positively correlated with both healthy and infected macrophage counts, and intracellular bacteria counts (**Figure 7a**). This relationship is expected because the initial number of macrophages (*N_itlMac*) determines the number of macrophages available to become infected and to phagocytose bacteria. Correspondingly, the initial macrophage count is negatively correlated with both planktonic and sessile bacteria counts. After the first 2-3 days of the simulation, the correlation between initial macrophage number and infected macrophage counts becomes negative, due to the decrease in infected macrophages from macrophage deaths.

The lasting correlations between initial macrophages and bacterial counts are because bacteria that are phagocytosed early in the simulation often stay intracellular for the remainder of the simulation. As we demonstrated above (Section 3.3), extracellular bacteria are more often a result of persistent extracellular bacteria, rather than those that were phagocytosed and released. Additionally, if left extracellular, a single planktonic bacterium can replicate between eight and sixteen times, yielding a potential of 256-65,000 extracellular bacteria over 14 days. In contrast, intracellular or sessile bacteria can only replicate 2-3 times or 4-8 times, respectively, yielding 8-256 bacteria over 14 days. In short, the high initial macrophage counts correlates negatively with planktonic, sessile, and eventually intracellular bacterial counts because the bacteria lost the opportunity to replicate quickly extracellularly, yielding fewer total bacteria over the course of the simulation.

Similarly, Initial Bacteria Count (*N_itlBac*) is positively correlated with infected macrophages and bacteria of all phenotypes throughout the simulation (**Figure 7b**). High initial bacterial counts also have a sustained negative correlation with healthy macrophage counts and a positive correlation with infected macrophages, because more macrophages are becoming infected. Thus, parameters that increase extracellular bacteria counts early in the simulation leads to higher bacterial levels longer-term. The fraction of initial bacteria that are planktonic (*F_itlPlank*) instead has a long-term impact on sessile bacteria, but not planktonic bacteria (**Figure 7c**). This observation leads to the question of how much bacterial growth rates impact bacterial numbers.

The bacterial doubling time parameter for planktonic (*T_plankBacDouble*) is negatively correlated with planktonic, sessile and intracellular bacteria counts (**Figure 7d**), while intracellular bacteria doubling time (*T_ICBacDouble*) is negatively correlated with intracellular bacteria counts (**Supplement 6aa**). In contrast, sessile doubling time (*T_sessBacDouble*) is not significantly correlated with any agent count (**Supplement 6f**). This contrast suggests that the sessile bacteria population is not sustained by replication, and that other sources such as initial deposition and phenotype switching play a more important role in sustaining sessile bacteria populations.

Finally, the *A_itlBiofilm* has a significant positive correlation with sessile bacterial counts, and a significant negative correlation with the number of macrophages, infected macrophages, and intracellular bacteria counts (**Figure 7e**). This correlation is in sharp contrast to *A_bacToBiofilm*, which is only significantly correlated for short periods (**Figure 7f**). This difference in impact of initial versus newly produced biofilm is due to the biological limits of the rate at which NTMs can make biofilm. A heatmap of biofilm levels at the final timepoint (**Figure 7g**) shows a gradient in total biofilm as a function of the initial amount of biofilm deposited with each sessile bacterium. However, within the biological range of new biofilm production by bacteria in the airways, there is almost no gradient in total biofilm levels. To confirm that this observation is due to the location within the parameter space, we completed another 900 simulations, gradually increasing the amount of biofilm above the biological range. It was not until each bacterium generated 1000-times the maximum of our biological range that bacterial contributions to new biofilm made a significant difference in the average amount of biofilm per grid square.

Overall, these findings show that initial parameters such as the initial number of bacteria and macrophages, bacterial phenotypic distributions, and biofilm (*N_itlBac*, *F_itlPlank*, *A_itlBiofilm*, *N_itlMac*) affect the long-term infection progression. These parameters do so either by changing the ratio of extracellular bacteria to macrophages early in the simulation (*N_itlBac* vs *N_itlMac*), or by increasing biofilm for protecting sessile bacteria (*A_itlBiofilm*). This preservation of extracellular bacteria forces the simulation down one of two pathways (predominantly intra- or extra-cellular), and thus ensures that those parameters remained significant throughout the simulation. Thus, initial parameters appear to determine the phenotype distribution in the simulation, and this phenotype distribution in turn determines the effective growth rate of the population, which drives overall bacterial numbers in the airways following MAC exposure.

## 4. Discussion

### 4.1 Establishing an infection

In this work, we use a computational model to explore the range of possible MAC infection outcomes focusing on early events before the recruitment of the adaptive immune system. These outcomes include complete bacterial clearance, all intracellular bacteria, and mixtures of intracellular, extracellular sessile and planktonic bacteria, and variations in macrophage recruitment.

#### 4.1.1 Initial parameters

We identify several key initial parameters (*N_itlBac, F_itlPlank*, and *A_itlBiofilm*) and showed how their effects propagate throughout the simulation. However, the true values or ranges for initial bacteria counts, phenotypes, or how much biofilm was present at the beginning remains unclear. Research has shown some correlations between environmental MAC sources and infections^6,61^. Further research into the phenotypic qualities of the bacteria that initially colonize the airway would further narrow our parameter ranges and add greatly to our understanding of how initial events impact MAC infections, and how they may be prevented.

Further, the significance of these initial parameters (*N_itlBac*, *N_itlMac*, *F_itlPlank*, and *A_itlBiofilm*) indicates that the course of infection may be decided early the airway colonization phase. Thus, development of a prophylactic treatment to prevent recurring infections^62^ may be an effective strategy. It is also possible that targeting these early pathogenic factors could also limit the progression of already established infections in the surrounding unaffected airways. This targeting may include either biofilms (which we discuss further below), stimulating mucosal clearance, and/or innate immune responses. Mucosal clearance is purposely neglected in this version of our model because individuals most at risk for NTM infections often have underlying conditions or structural lung disease such as bronchiectasis or chronic obstructive pulmonary disease that impair mucous clearance.

#### 4.1.2 Macrophage Response

Our findings regarding the importance of balancing macrophage recruitment is consistent with tuberculosis where the spectrum of insufficient recruitment to high levels of recruitment and inflammation impacts disease progression and granuloma formation^63^. The fragility of this balance indicates that directly targeting macrophage recruitment, either to stimulate or suppress their response, would not be an effective way to prevent NTM-PD. Instead, these results have increased our interest in the role of biofilms in increasing inflammation via apoptosis, and the effects of biofilms across diverse macrophage phenotypes and other components of the early immune response.

A key assumption of this model is that there is a dose-response relationship between amount of biofilm and its effect on macrophages, both inhibition of phagocytosis and apoptosis. This assumption could be improved with *in vitro* experiments that measure the interactions between macrophages, bacteria and biofilms, especially the distinction between microaggregates (precursors to sessile bacteria clusters) and sessile bacteria^20,22^. We know that microaggregates have increased probabilities of being phagocytosed^22^, but that bacteria in mature biofilms are less likely to be phagocytosed compared to planktonic bacteria^19^. Further elucidation of the role of progressively increasing levels of biofilm in phagocytosis will allow us to refine our dose-response assumption.

We have also seen that macrophage deaths are largely caused by these biofilms, for which we assume a linear dose-response relationship, but experimental studies have only shown an effect with mature biofilm (Rose & Bermudez, 2014). Better understanding this relationship would allow us to further probe potential effects potential prophylactic and therapeutic interventions via methods such as partially degrading biofilms or attenuating macrophages’ apoptotic response to them.

### 4.2 Implications of diverse model endpoints for infection type

Variations in bacterial phenotypic distribution may explain the lack of correlation of treatment efficacy with *in vitro* minimum inhibitory concentrations (MIC) measurements, if bacteria being produced in sputum samples are not representative of largely-intracellular populations. Some studies have examined the differences of MICs in bacteria alone or within macrophage cultures, and have found that the MICs and minimal bactericidal concentrations (MBCs) of intra- and extracellular bacteria are different, but neither one is consistently lower than the other across the different antibiotics tested^64^. This discrepancy indicates that understanding the distribution of bacterial phenotypes and locations (intra- vs. extracellular) in the different stages of MAC pulmonary infection can have significant clinical implications for therapeutic interventions.

Despite the diversity of predominant phenotypes (intracellular, planktonic or sessile) and bacterial loads, the likelihood of each phenotype being dominant is not equal. We see a majority of bacteria were intracellular at the end of the 14-day simulation, indicating that bacteria in an infection will be predominantly within macrophages. The activation of the immune system at the end of the 14 days post-colonization could cause many of the intracellular bacteria to be cleared. There are, however, two scenarios of all-intracellular bacteria that may inhibit this clearance. First if all bacteria are intracellular but macrophages have not been able to recruit any other immune cells, induction of the adaptive immune system may be delayed, requiring macrophages to kill intracellular bacteria without activation, or eventually burst. Second if infected macrophages recruited many healthy macrophages, we may see barrier effects where recruited cytotoxic T cells fail to reach bacteria, preventing activation of infected macrophages.

If infected macrophages cannot be activated, they can cause further infection in two ways. First, they may eventually burst releasing bacteria back into the mucosal environment, which will allow bacteria to grow extracellularly in the mucus layer as seen in other simulations with predominantly extracellular bacteria. Otherwise, infected macrophages may move deeper into lung tissue^65^ eventually forming infections similar to those seen in tuberculosis infections and nodular cases in NTM-PD.

Meanwhile, extracellular planktonic and sessile bacteria do not necessarily mean an infection will progress, as bacteria in the airway are still only considered colonization, rather than infection. The larger threat that these extracellular bacteria present is sessile bacteria in biofilm invading epithelial cells.

This difference leads to broader implications of these early colonization and infection events for the course of an infection. We know that nodular and cavitary MAC-lung disease have different outcomes and treatment responses to antimicrobials. It has been suggested that a combination of host immune and bacterial factors may account for these two different disease presentations^66^. Based on the literature and our findings, we hypothesize that the bacterial populations with relatively high sessile bacteria counts may eventually lead to cavitary disease, while higher levels of intracellular bacteria may lead to nodular disease as infection progresses. This theory is supported by the following: First, we see that higher levels of initial biofilm combined with higher initial sessile bacteria counts correlates well with more sessile bacteria and more invaded bacteria over time. These invaded bacteria will have a biofilm-producing phenotype. Bacterial biofilms are also well-documented to attenuate antibiotic effects^7,67^, which may partially explain cavitary infections’ resistance to treatment. Second, we know that *in vitro* antibiotic susceptibility tests have a low correlation with treatment efficacy^68,69^. Here we have shown a variety of bacterial phenotypes present at the end of simulations. Bacteria remaining in the airway will not necessarily have the same phenotype as those that invaded. We believe that these airway bacteria most closely represent what would be found in sputum samples, which may explain the lack of correlation seen in clinical settings. Finally, fibrocavitary disease usually presents in the upper lobe of lungs^70^. Studies have shown that particles as large as sessile bacteria encased in biofilm are more likely to experience impaction into mucus earlier after inhalation, and are therefore deposited higher in the airways^71^, and more biofilm surrounding an inhaled sessile bacterium will increase its size. Therefore, higher initial biofilm will increase the likelihood of a bacterium with biofilm being deposited in upper zone branches of the lungs^72^.

Understanding how different disease presentations occur will aid development of both treatment and prevention strategies.

### 4.3 Model Limitations

While this model probes factors that lead to a variety of possible outcomes of MAC colonization of the lungs, it has limitations. As discussed, patients who are susceptible to NTM infections often have pre-existing factors such as COPD, cystic fibrosis, or bronchiectasis which affects the lung airways. While we purposely neglect mucosal clearance because of these factors, we did not examine the role of pre-existing inflammation or damage in the airways, and they will vary from patient to patient depending on airway microbiota and past conditions. Correspondingly, there are various therapies used by patients with these conditions for airway health (airway clearance techniques, nebulized hypertonic saline, bronchodilators, corticosteroid therapies, etc.) that will also impact immune response and bacterial clearance.

These considerations, in combination with parameterization limitations (initial bacterial inoculum, apoptotic response to biofilms, etc.) mean that while our model can explore and explain some of the effects that are seen in the lungs, it cannot not make individual patient predictions. Future work, however, will address these concerns and include patient-specific information to test our ability to make predictions for clinical care.

## 5. Conclusions

We present an agent-based model of the mucosal layer of the lung airway populated with MAC bacteria and host macrophages in order to explore the interactions between bacteria, their biofilms, and host macrophages during the initiation of NTM-PD (**Figure 1**). These behaviors and dynamics are integrated from diverse but isolated experiments and combined to provide an understanding of this complex system.

We predict that initial conditions, host immune response, and bacterial biofilm play large roles in deciding the outcome of exposure leading to either bacterial clearance, continued colonization, or the initiation of an infection. We believe that these diverse outcomes may in part explain the different presentations of MAC lung disease and other NTM-PD, either fibrocavitary or nodular bronchiectatic disease. Further exploration of biofilm-macrophage interactions in this model may indicate novel drug targets and identify potential new preventive and therapeutic strategies for patients at risk of new and recurrent NTM pulmonary infections.

## 6. Acknowledgements

This research was supported by a grant from the CHEST Foundation in partnership with Insmed Incorporated (E.P. and P.E.); and the Frederick N. Andrews Fellowship (C.W.). This paper’s contents are solely the responsibility of the authors and do not necessarily represent the official views of the CHEST Foundation, Mayo Clinic, or any other organization. We thank Jonathan Ozik, Eric Tatara, and Nick Collier for their expertise and support with Repast Simphony. We also thank Lev Gorenstein, Tsai-Wei Wu, and the rest of the Research Computing Staff for their assistance with batch computing at the Rosen Center for Advanced Computing.

## References

1. Johnson, M. M. & Odell, J. A. Nontuberculous mycobacterial pulmonary infections. J. Thorac. Dis. 6, 210–220 (2014).

2. Carter, G., Wu, M., Drummond, D. C. & Bermudez, L. E. Characterization of biofilm formation by clinical isolates of Mycobacterium avium. J. Med. Microbiol. 52, 747–752 (2003).

3. Cassidy, P. M., Hedberg, K., Saulson, A., McNelly, E. & Winthrop, K. L. Nontuberculous Mycobacterial Disease Prevalence and Risk Factors: A Changing Epidemiology. Clin. Infect. Dis. 49, e124–e129 (2009).

4. van Ingen, J., Ferro, B. E., Hoefsloot, W., Boeree, M. J. & van Soolingen, D. Drug treatment of pulmonary nontuberculous mycobacterial disease in HIV-negative patients: the evidence. Expert Rev. Anti Infect. Ther. 11, 1065–1077 (2013).

5. Nishiuchi, Y., Iwamoto, T. & Maruyama, F. Infection Sources of a Common Non-tuberculous Mycobacterial Pathogen, Mycobacterium avium Complex. Front. Med. 4, (2017).

6. Tzou, C. L. et al. Association between *Mycobacterium avium* Complex Pulmonary Disease and Mycobacteria in Home Water and Soil: A Case–Control Study. Ann. Am. Thorac. Soc. 17, 57–62 (2020).

7. Falkinham, J. O. Growth in catheter biofilms and antibiotic resistance of Mycobacterium avium. J. Med. Microbiol. 56, 250–254 (2007).

8. Falkinham, J. O. Disinfection and cleaning of heater–cooler units: suspension- and biofilm-killing. J. Hosp. Infect. 105, 552–557 (2020).

9. Flemming, H.-C. et al. Biofilms: an emergent form of bacterial life. Nat. Rev. Microbiol. 14, 563–575 (2016).

10. Daniel-Wayman, S. et al. Advancing Translational Science for Pulmonary Nontuberculous Mycobacterial Infections. A Road Map for Research. Am. J. Respir. Crit. Care Med. 199, 947–951 (2019).

11. Fennelly, K. P. et al. Biofilm Formation by *Mycobacterium abscessus* in a Lung Cavity. Am. J. Respir. Crit. Care Med. 193, 692–693 (2016).

12. Qvist, T. et al. Chronic pulmonary disease with *Mycobacterium abscessus* complex is a biofilm infection. Eur. Respir. J. 46, 1823–1826 (2015).

13. Chakraborty, P., Bajeli, S., Kaushal, D., Radotra, B. D. & Kumar, A. Biofilm formation in the lung contributes to virulence and drug tolerance of Mycobacterium tuberculosis. Nat. Commun. 12, 1606 (2021).

14. Brown-Elliott, B. A., Nash, K. A. & Wallace, R. J. J. Antimicrobial susceptibility testing, drug resistance mechanisms, and therapy of infections with nontuberculous mycobacteria. Clin. Microbiol. Rev. 25, 545–582 (2012).

15. Esteban, J. & García-Coca, M. Mycobacterium Biofilms. Front. Microbiol. 8, 2651 (2018).

16. Lewis, A. H. & Falkinham, J. O. Microaerobic growth and anaerobic survival of Mycobacterium avium, Mycobacterium intracellulare and Mycobacterium scrofulaceum. Int. J. Mycobacteriology 4, 25–30 (2015).

17. Carter, G., Young, L. S. & Bermudez, L. E. A Subinhibitory Concentration of Clarithromycin Inhibits *Mycobacterium avium* Biofilm Formation. Antimicrob. Agents Chemother. 48, 4907–4910 (2004).

18. Williams, M. M. et al. Structural Analysis of Biofilm Formation by Rapidly and Slowly Growing Nontuberculous Mycobacteria. Appl. Environ. Microbiol. 75, 2091–2098 (2009).

19. Rose, S. J. & Bermudez, L. E. Mycobacterium avium Biofilm Attenuates Mononuclear Phagocyte Function by Triggering Hyperstimulation and Apoptosis during Early Infection. Infect. Immun. 82, 405–412 (2014).

20. Yamazaki, Y. et al. The ability to form biofilm influences Mycobacterium avium invasion and translocation of bronchial epithelial cells. Cell. Microbiol. 8, 806–814 (2006).

21. Dominici, S., Brandi, G., Schiavano, G. F. & Magnani, M. Selective Killing of Mycobacterium avium–Infected Macrophages by Inhibition of Phosphorylated Signal Transducer and Activator of Transcription Type 1. J. Infect. Dis. 198, 95–100 (2008).

22. Keefe, B. F., Leestemaker-Palmer, A. & Bermudez, L. E. Mycobacterium avium subsp. hominissuis (MAH) Microaggregate induction of host innate immunity is linked to biofilm formation. Microb. Pathog. 157, 104977 (2021).

23. Ratnatunga, C. N. et al. The Rise of Non-Tuberculosis Mycobacterial Lung Disease. Front. Immunol. 11, 303 (2020).

24. Koh, W.-J. et al. Outcomes of *Mycobacterium avium* complex lung disease based on clinical phenotype. Eur. Respir. J. 50, 1602503 (2017).

25. Hwang, J. A., Kim, S., Jo, K.-W. & Shim, T. S. Natural history of *Mycobacterium avium* complex lung disease in untreated patients with stable course. Eur. Respir. J. 49, 1600537 (2017).

26. Griffith, D. E. et al. An Official ATS/IDSA Statement: Diagnosis, Treatment, and Prevention of Nontuberculous Mycobacterial Diseases. Am. J. Respir. Crit. Care Med. 175, 367–416 (2007).

27. Orme, I. M. & Ordway, D. J. Host Response to Nontuberculous Mycobacterial Infections of Current Clinical Importance. Infect. Immun. 82, 3516–3522 (2014).

28. Cadena, A. M., Flynn, J. L. & Fortune, S. M. The Importance of First Impressions: Early Events in Mycobacterium tuberculosis Infection Influence Outcome. mBio 7, (2016).

29. Davis, J. M. & Ramakrishnan, L. The Role of the Granuloma in Expansion and Dissemination of Early Tuberculous Infection. Cell 136, 37–49 (2009).

30. Redford, P. S. et al. Enhanced protection to Mycobacterium tuberculosis infection in IL-10-deficient mice is accompanied by early and enhanced Th1 responses in the lung. Eur. J. Immunol. 40, 2200–2210 (2010).

31. Bermudez, L. E. et al. Activity of Moxifloxacin by Itself and in Combination with Ethambutol, Rifabutin, and Azithromycin In Vitro and In Vivo against *Mycobacterium avium*. Antimicrob. Agents Chemother. 45, 217–222 (2001).

32. Gangadharam, P. R. J. et al. Susceptibility of Beige Mice to *Mycobacterium avium* Complex Infections by Different Routes of Challenge. Am. Rev. Respir. Dis. 139, 1098–1104 (1989).

33. Andréjak, C. et al. Characterization of Mouse Models of Mycobacterium avium Complex Infection and Evaluation of Drug Combinations. Antimicrob. Agents Chemother. 59, 2129–2135 (2015).

34. Bonabeau, E. Agent-based modeling: Methods and techniques for simulating human systems. Proc. Natl. Acad. Sci. 99, 7280–7287 (2002).

35. Kirschner, D., Pienaar, E., Marino, S. & Linderman, J. J. A review of computational and mathematical modeling contributions to our understanding of Mycobacterium tuberculosis within-host infection and treatment. Curr. Opin. Syst. Biol. 3, 170–185 (2017).

36. Gopalakrishnan, V., Kim, M. & An, G. Using an Agent-Based Model to Examine the Role of Dynamic Bacterial Virulence Potential in the Pathogenesis of Surgical Site Infection. Adv. Wound Care 2, 510–526 (2013).

37. Chambless, J. D. & Stewart, P. S. A three-dimensional computer model analysis of three hypothetical biofilm detachment mechanisms. Biotechnol. Bioeng. 97, 1573–1584 (2007).

38. North, M. J. et al. Complex adaptive systems modeling with Repast Simphony. Complex Adapt. Syst. Model. 1, 3 (2013).

39. Lindestam Arlehamn, C. S., Lewinsohn, D., Sette, A. & Lewinsohn, D. Antigens for CD4 and CD8 T Cells in Tuberculosis. Cold Spring Harb. Perspect. Med. 4, a018465–a018465 (2014).

40. Wolf, A. J. et al. Initiation of the adaptive immune response to Mycobacterium tuberculosis depends on antigen production in the local lymph node, not the lungs. J. Exp. Med. 205, 105–115 (2008).

41. Ichiyama, S., Shimokata, K. & Tsukamura, M. The Isolation of *Mycobacterium avium* Complex from Soil, Water, and Dusts. Microbiol. Immunol. 32, 733–739 (1988).

42. Kirschner, R. A., Parker, B. C. & Falkinham, J. O. Epidemiology of Infection by Nontuberculous Mycobacteria: *Mycobacterium avium, Mycobacterium intracellulare*, and *Mycobacterium scrofulaceum* in Acid, Brown-Water Swamps of the Southeastern United States and Their Association with Environmental Variables. Am. Rev. Respir. Dis. 145, 271–275 (1992).

43. Falkinham, J. O., Norton, C. D. & LeChevallier, M. W. Factors Influencing Numbers of *Mycobacterium avium*, *Mycobacterium intracellulare*, and Other Mycobacteria in Drinking Water Distribution Systems. Appl. Environ. Microbiol. 67, 1225–1231 (2001).

44. Falkinham, J. O. Hospital water filters as a source of Mycobacterium avium complex. J. Med. Microbiol. 59, 1198–1202 (2010).

45. Glazer, C. S. et al. Nontuberculous Mycobacteria in Aerosol Droplets and Bulk Water Samples from Therapy Pools and Hot Tubs. J. Occup. Environ. Hyg. 4, 831–840 (2007).

46. Falkinham, J. O. Mycobacterial Aerosols and Respiratory Disease. Emerg. Infect. Dis. 9, 763–767 (2003).

47. Chen, M. J., Zhang, Z. & Bott, T. R. Direct measurement of the adhesive strength of biofilms in pipes by micromanipulation. Biotechnol. Tech. 12, 875–880 (1998).

48. Pienaar, E., Matern, W. M., Linderman, J. J., Bader, J. S. & Kirschner, D. E. Multiscale Model of Mycobacterium tuberculosis Infection Maps Metabolite and Gene Perturbations to Granuloma Sterilization Predictions. Infect. Immun. 84, 1650–1669 (2016).

49. Early, J., Fischer, K. & Bermudez, L. E. Mycobacterium avium uses apoptotic macrophages as tools for spreading. Microb. Pathog. 50, 132–139 (2011).

50. Cilfone, N. A., Kirschner, D. E. & Linderman, J. J. Strategies for Efficient Numerical Implementation of Hybrid Multi-scale Agent-Based Models to Describe Biological Systems. Cell. Mol. Bioeng. 8, 119–136 (2015).

51. Wallace, W. A., Gillooly, M. & Lamb, D. Intra-alveolar macrophage numbers in current smokers and non-smokers: a morphometric study of tissue sections. Thorax 47, 437–440 (1992).

52. Warheit, D. B. & Hartsky, M. A. Role of alveolar macrophage chemotaxis and phagocytosis in pulmonary clearance responses to inhaled particles: Comparisons among rodent species. Microsc. Res. Tech. 26, 412–422 (1993).

53. Harkness, L. M., Kanabar, V., Sharma, H. S., Westergren-Thorsson, G. & Larsson-Callerfelt, A.-K. Pulmonary vascular changes in asthma and COPD. Pulm. Pharmacol. Ther. 29, 144–155 (2014).

54. Lukacs, N. W., Strieter, R. M., Chensue, S. W., Widmer, M. & Kunkel, S. L. TNF-alpha mediates recruitment of neutrophils and eosinophils during airway inflammation. J. Immunol. Baltim. Md 1950 154, 5411–5417 (1995).

55. Sun, L. et al. Effect of IL-10 on Neutrophil Recruitment and Survival after *Pseudomonas aeruginosa* Challenge. Am. J. Respir. Cell Mol. Biol. 41, 76–84 (2009).

56. Hacker, T., Yang, B. & McCartney, G. Empowering Faculty: A Campus Cyberinfrastructure Strategy for Research Communities. Educ. Rev. (2014).

57. Rahaghi, F. N. et al. Pulmonary vascular density: comparison of findings on computed tomography imaging with histology. Eur. Respir. J. 54, 1900370 (2019).

58. Crowle, A. J., Tsang, A. Y., Vatter, A. E. & May, M. H. Comparison of 15 laboratory and patient-derived strains of Mycobacterium avium for ability to infect and multiply in cultured human macrophages. J. Clin. Microbiol. 24, 812–821 (1986).

59. Goodhill, G. J. Diffusion in Axon Guidance. Eur. J. Neurosci. 9, 1414–1421 (1997).

60. Serisier, D. J., Carroll, M. P., Shute, J. K. & Young, S. A. Macrorheology of cystic fibrosis, chronic obstructive pulmonary disease & normal sputum. Respir. Res. 10, 63 (2009).

61. Thomson, R. et al. Isolation of Nontuberculous Mycobacteria (NTM) from Household Water and Shower Aerosols in Patients with Pulmonary Disease Caused by NTM. J. Clin. Microbiol. 51, 3006–3011 (2013).

62. Boyle, D. P., Zembower, T. R. & Qi, C. Relapse versus Reinfection of *Mycobacterium avium* Complex Pulmonary Disease. Patient Characteristics and Macrolide Susceptibility. Ann. Am. Thorac. Soc. 13, 1956–1961 (2016).

63. Roach, D. R. et al. TNF Regulates Chemokine Induction Essential for Cell Recruitment, Granuloma Formation, and Clearance of Mycobacterial Infection. J. Immunol. 168, 4620–4627 (2002).

64. Chanwong, S., Maneekarn, N., Makonkawkeyoon, L. & Makonkawkeyoon, S. Intracellular growth and drug susceptibility of Mycobacterium tuberculosis in macrophages. Tuberculosis 87, 130–133 (2007).

65. Cohen, S. B. et al. Alveolar Macrophages Provide an Early Mycobacterium tuberculosis Niche and Initiate Dissemination. Cell Host Microbe 24, 439–446.e4 (2018).

66. Abe, Y. et al. Host Immune Response and Novel Diagnostic Approach to NTM Infections. Int. J. Mol. Sci. 21, 4351 (2020).

67. Rose, S. J., Babrak, L. M. & Bermudez, L. E. Mycobacterium avium Possesses Extracellular DNA that Contributes to Biofilm Formation, Structural Integrity, and Tolerance to Antibiotics. PLOS ONE 10, e0128772 (2015).

68. Heginbothom, M. L. The relationship between the in vitro drug susceptibility of opportunist mycobacteria and their in vivo response to treatment. Int. J. Tuberc. Lung Dis. Off. J. Int. Union Tuberc. Lung Dis. 5, 539–545 (2001).

69. Kobashi, Y., Yoshida, K., Miyashita, N., Niki, Y. & Oka, M. Relationship between clinical efficacy of treatment of pulmonary Mycobacterium avium complex disease and drugsensitivity testing of Mycobacterium avium complex isolates. J. Infect. Chemother. Off. J. Jpn. Soc. Chemother. 12, 195–202 (2006).

70. Yoo, S. H. et al. Multiple Cavitary Pulmonary Nodules Caused by *Mycobacterium intracellulare*. Korean J. Fam. Med. 37, 248 (2016).

71. Heyder, J. Deposition of Inhaled Particles in the Human Respiratory Tract and Consequences for Regional Targeting in Respiratory Drug Delivery. Proc. Am. Thorac. Soc. 1, 315–320 (2004).

72. Frederix, E. M. A. et al. Simulation of size-dependent aerosol deposition in a realistic model of the upper human airways. J. Aerosol Sci. 115, 29–45 (2018).

