## Supplements for "The Role of Biofilms, Bacterial Phenotypes, and Innate Immune Response in *Mycobacterium avium* Colonization to Infection"

### Supplement 1

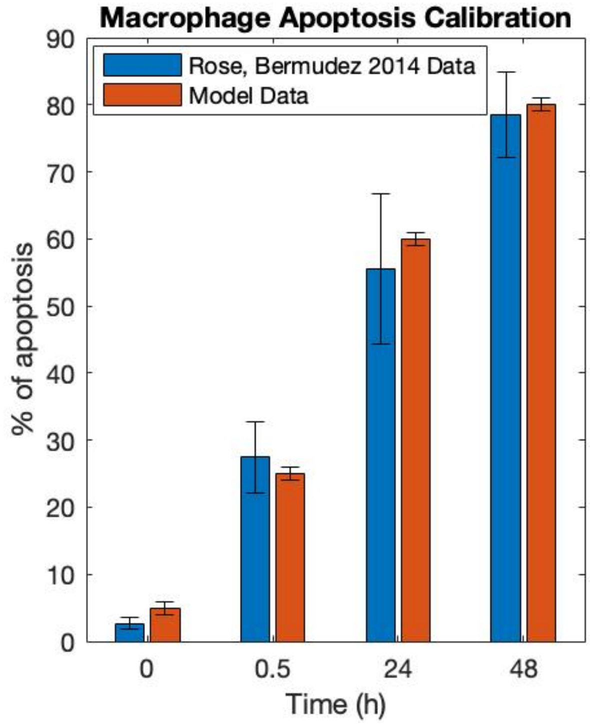


Calibration of Macrophage Apoptosis in Response to Biofilm Exposure. Macrophage tolerance of biofilms is calibrated to data from Rose & Bermudez 2014[19], which added THP-1 cells to fully formed biofilms and counted apoptotic cells. To calibrate, a simulation with 100% biofilm is created and macrophages were added. Over time apoptosis is counted. Proportions of apoptosed cells in the published Rose data and model are statistically identical at each timepoint (α=0.05).

### Supplement 2

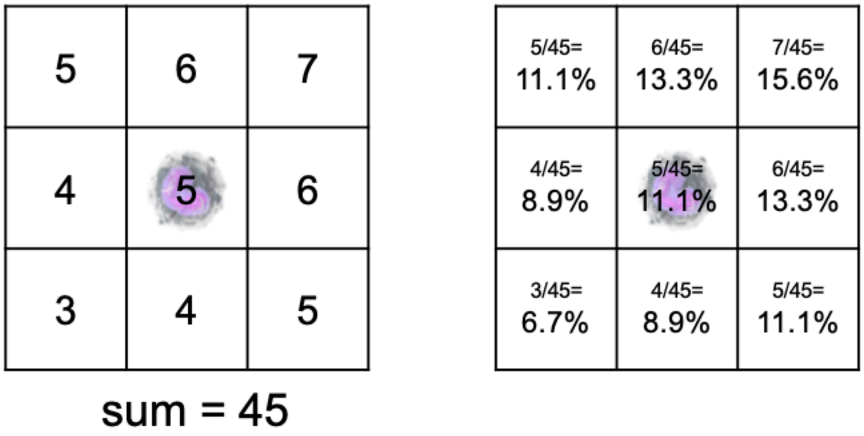


Macrophage Chemotaxis Algorithm. Macrophages undergo chemotaxis, probabilistically moving up the chemoattractant gradient, by weighting the probability of moving to a gridsquare in its Moore neighborhood by chemoattractant values. In this diagram, the macrophage (blue) is at the center and examines its Moore neighborhood for chemoattractant values (left), then calculates the probability of chemotaxing to each (right). Direction is then chosen randomly with each gridsquare weighted correspondingly.

### Supplement 3

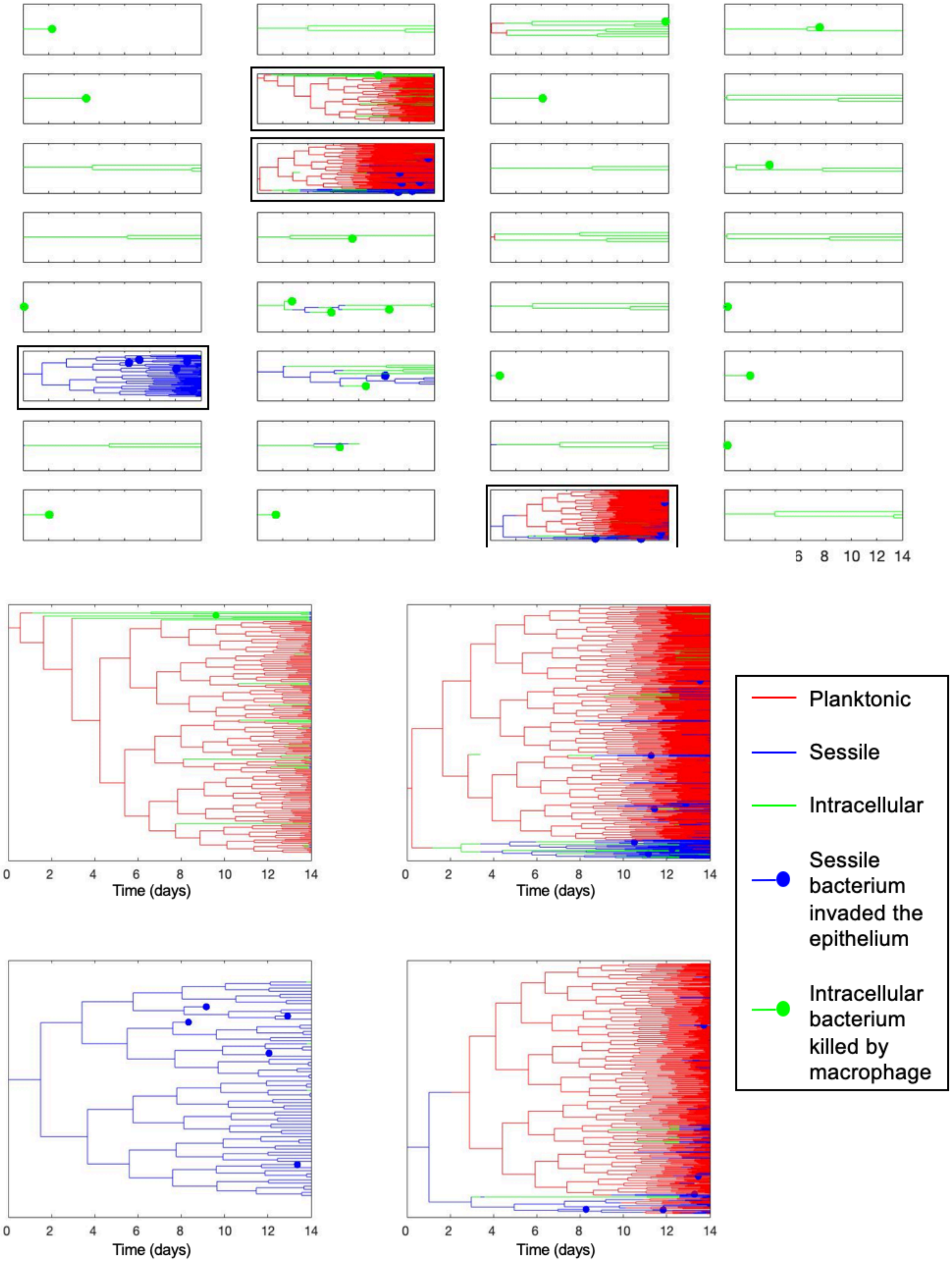


Bacteria Dynamics in a representative simulation. Each small graph represents a single initial bacterium and all of its descendants, color-coded for their phenotype. Dots at the end of the line indicate that the bacterium was either killed (it it was green, for intracellular) or invaded the epithelium (blue, for sessile), and was no longer in the simulation. The four graphs in boxes are shown in greater detail below, so that we can more easily see the dynamics of the bacteria.

**Supplement 4**

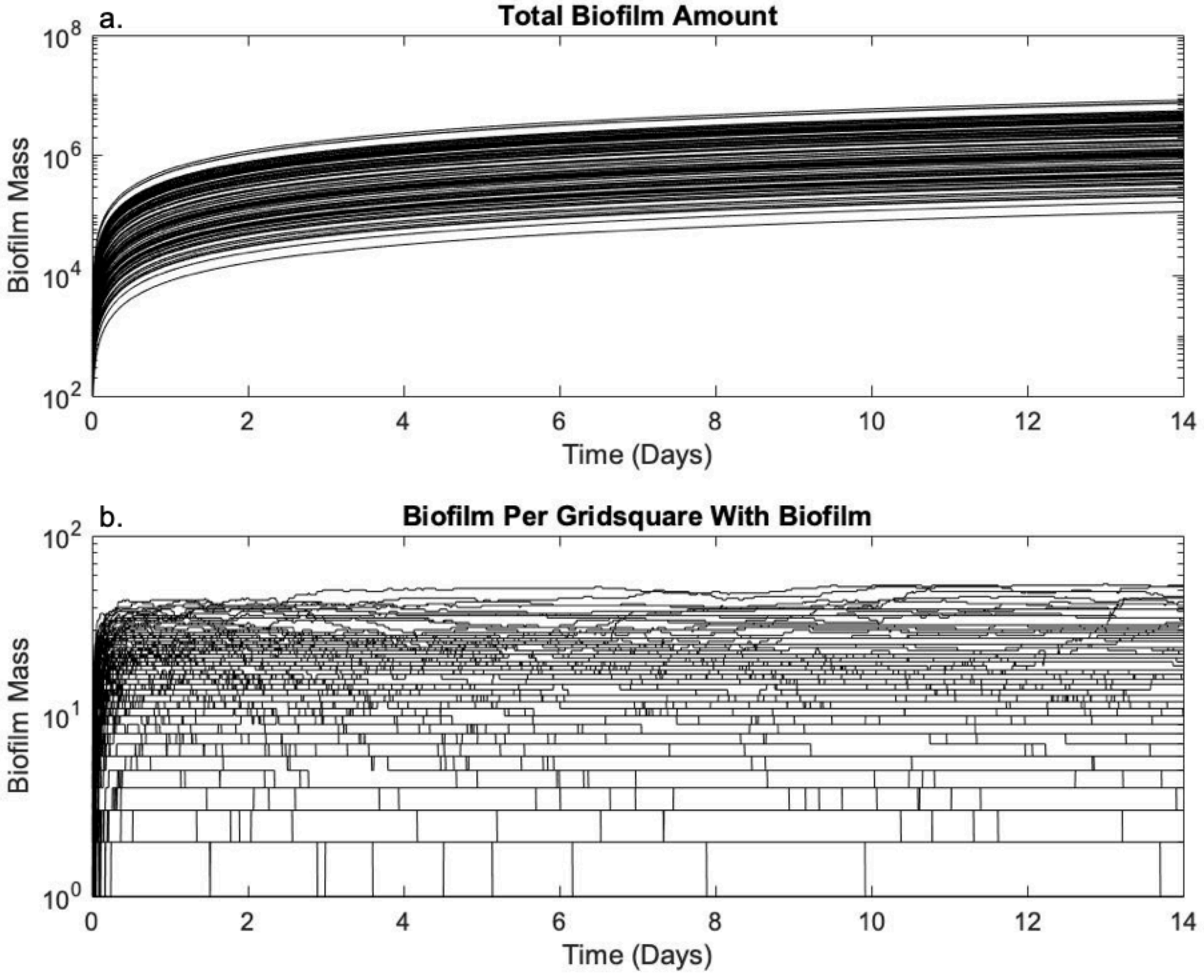


Biofilm amounts over time. a) shows the total mass of biofilm in the airway over time, while b) shows the amount of biofilm per gridsquare that had biofilm, indicating the amount of protection for local sessile bacteria.

### Supplement 5

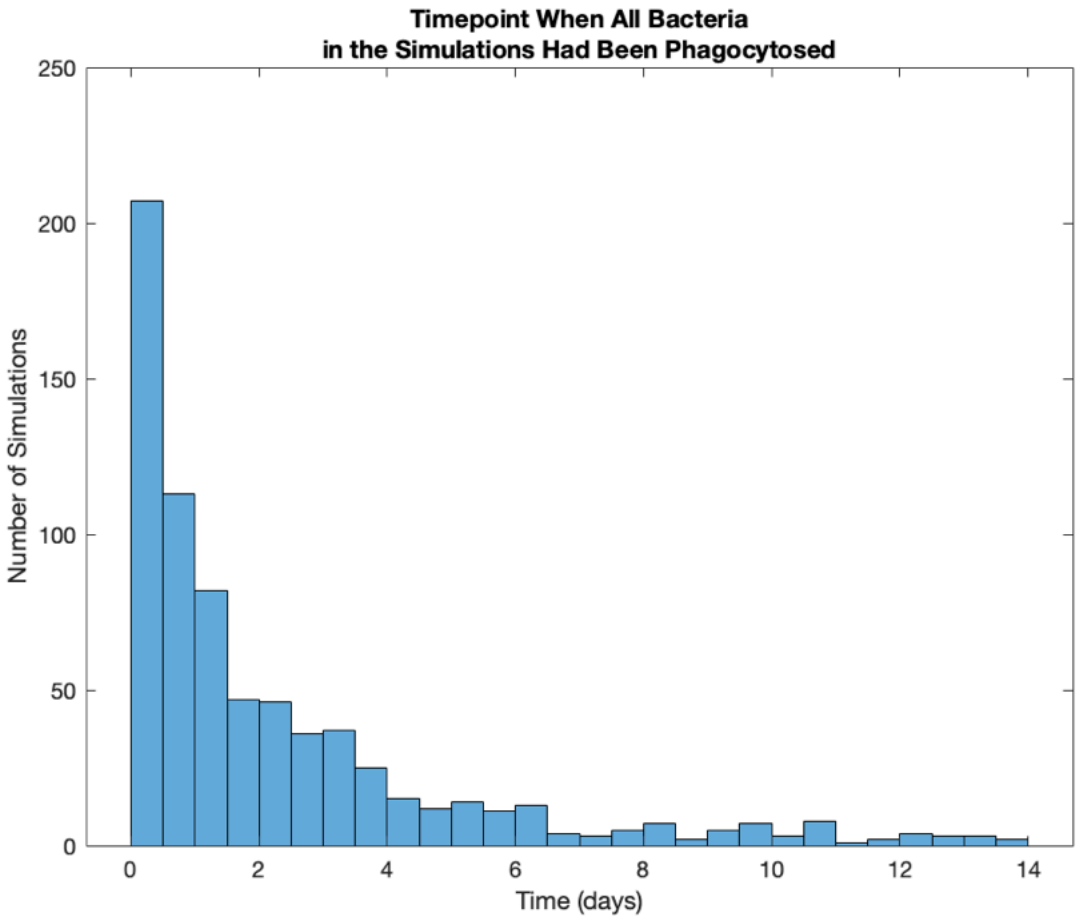


Histogram of the timepoint when all bacteria became intracellular for each simulation.

**Supplement 6**

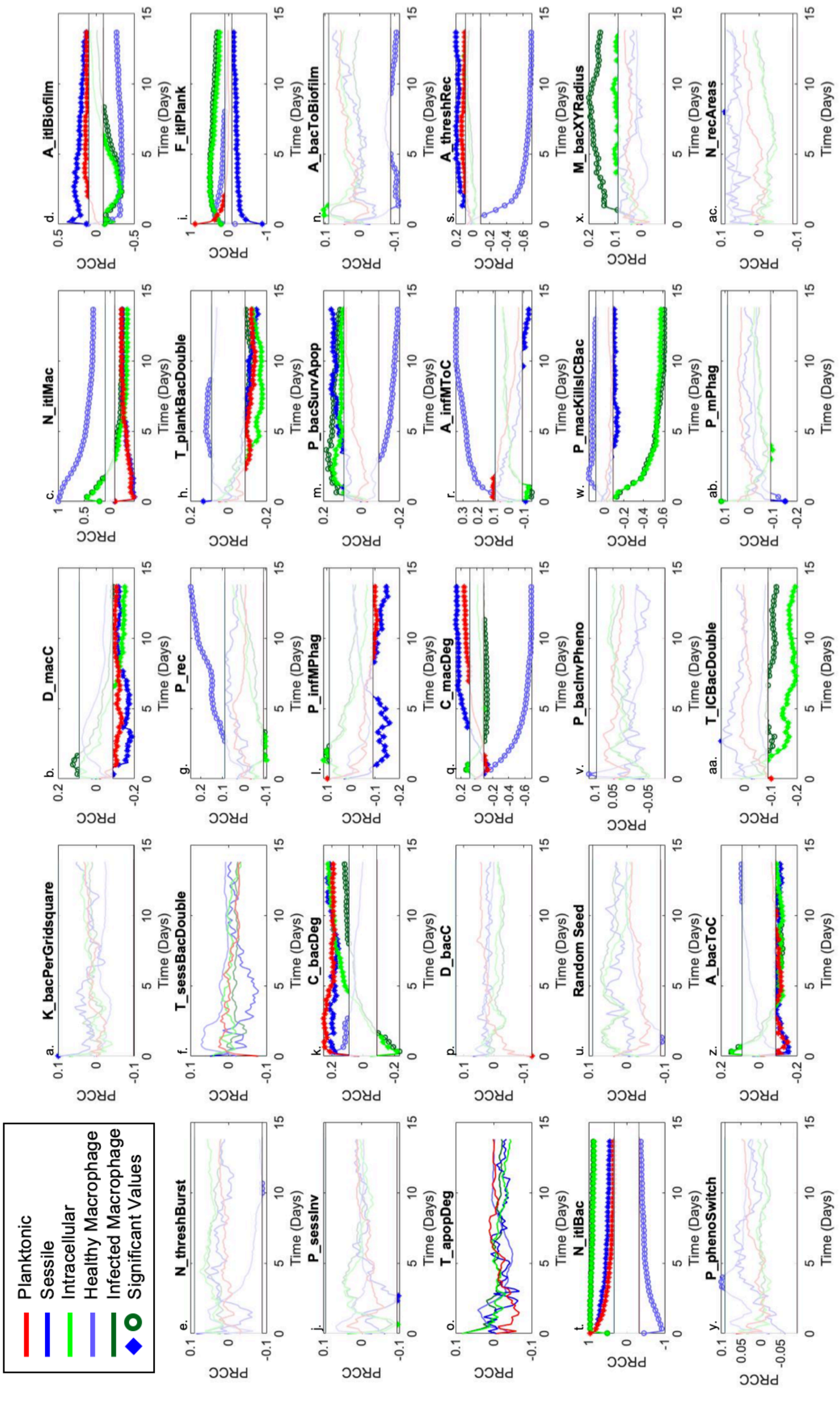

Partial Rank Correlation Coefficient of each parameter on planktonic, sessile, and intracellular bacterial counts, and healthy and infected macrophage counts. The range in which these values are not significant centers around 0, and the threshold is marked by a horizontal black line above and below. Values in the insignificant range are plotted as lines, but do not have individual data markers and are greyed out. Significant values are marked, using a filled point for bacteria or an open point for macrophages. In each plot, intracellular and infected macrophage coe
